# Distinct and shared synergistic effects of high-fat and high-iron diets on liver tumorigenesis and transcriptomic remodeling in mice

**DOI:** 10.64898/2026.05.24.727480

**Authors:** Debodipta Das, Hakim Bouamar, Xiaoli Sun, Jason Xu, Lu Cai, Yidong Chen, Francis E. Sharkey, Sukeshi P. Arora, Francisco G. Cigarroa, Lu-Zhe Sun

**Affiliations:** Department of Cell Systems & Anatomy, University of Texas Health Science Center at San Antonio, TX 78229, USA; Department of Pharmacology, University of Texas Health Science Center at San Antonio, TX 78229, USA; Pediatric Research Institute, Department of Pediatrics, University of Louisville School of Medicine, Louisville KY 40202, USA; Greehey Children’s Cancer Research Institute, University of Texas Health Science Center at San Antonio, TX 78229, USA; Department of Population Health Sciences, University of Texas Health Science Center at San Antonio, TX 78229, USA; Department of Pathology and Laboratory Medicine, University of Texas Health Science Center at San Antonio, TX 78229, USA; Department of Medicine, University of Texas Health Science Center at San Antonio, TX 78229, USA; Transplant Center, University of Texas Health Science Center at San Antonio, TX 78229, USA; Centre for Integrative Omics Data Science (CIODS), Yenepoya (Deemed to be University), Mangalore 575018, INDIA

**Keywords:** Hepatocellular carcinoma, High-fat diet, Dietary iron overload, mTOR signaling, Diet-induced transcriptomic reprogramming

## Abstract

**Background and Aims:** The incidence of hepatocellular carcinoma (HCC) is increasingly driven by metabolic risk factors, including obesity and iron overload. Although high-fat and high-iron diets independently promote hepatocarcinogenesis, their shared and distinct molecular effects remain unclear. We sought to define how dietary fat and iron differentially shape HCC development.

**Approach:** Male C3HeB/FeJ mice were exposed to long-term high-fat (HFD), high-iron (HID), or combined (HFD+HID) diets for 16.5 months. Tumor burden, hepatic iron distribution, mTOR signaling, oxidized phospholipid (OxPL) accumulation, and transcriptomic alterations across normal, adjacent non-tumor, and tumor liver tissues were analyzed using biochemical, histological, and RNA sequencing approaches.

**Results:** All diets induced HCC with comparable tumor burden. HID increased iron levels in non-tumor liver tissue but resulted in relative iron depletion within tumors, indicating tumor-specific iron utilization. Tumors from all diet groups showed robust mTOR activation and increased OxPL accumulation, with stronger oxidative stress signatures in HFD and HFD+HID tumors. Transcriptomic analyses revealed conserved oncogenic programs alongside diet-specific signatures, with HFD exerting a dominant effect on metabolic reprogramming and gene dysregulation, whereas HID preferentially enhanced immune and inflammatory signaling. Progressive, monotonic changes in gene expression were observed across disease stages. Cross-species analyses linked diet-induced mouse tumors to immunologically “hot” human HCC subtypes.

**Conclusions:** Dietary fat and iron promote HCC through overlapping yet distinct molecular pathways, highlighting metabolic and immune mechanisms as key targets in diet-associated liver cancer.

## Introduction

Hepatocellular carcinoma (HCC), the most common form of primary liver cancer, is a major contributor to global cancer-related morbidity and mortality (1). While chronic infections with hepatitis B and C viruses remain the leading global risk factors, metabolic conditions such as obesity, metabolic dysfunction-associated steatotic liver disease (MASLD), and iron overload disorders are emerging as significant contributors to the increasing burden of HCC, especially in high-income and transitioning countries (2–5).

Dietary factors, particularly high-fat and high-iron intake, have been implicated in liver pathophysiology and tumorigenesis through mechanisms involving metabolic reprogramming, inflammation, oxidative stress, and aberrant signaling pathway activation (6–8). High-fat diets (HFD) are known to promote obesity-associated liver steatosis and inflammation, which can progress to metabolic dysfunction-associated steatohepatitis (MASH), fibrosis, and eventually HCC (9,10). Similarly, excess dietary iron can lead to hepatic iron accumulation and oxidative damage through Fenton chemistry, resulting in hepatocyte injury, inflammation, and neoplastic transformation (11,12).

Despite this knowledge, the interplay between dietary fat and iron in driving hepatocarcinogenesis remains poorly understood. Previous studies have typically examined the effects of either high-fat (13) or high-iron (14) diets in isolation, with limited insight into their combined impact on tumor biology and molecular pathways (8). Moreover, while obesity and iron overload are individually associated with activation of oncogenic signaling cascades such as mammalian target of rapamycin (mTOR) and increased lipid peroxidation, their integrated effects on liver tissue remodeling, tumor microenvironment, and gene expression landscapes have not been systematically investigated.

The C3HeB/FeJ mouse strain provides a spontaneous, non-viral model of hepatocarcinogenesis that closely mirrors key aspects of human disease, including male predominance, middle-age onset, and histopathological features such as steatosis, tumor progression, and vascular invasion (15). With a ∼50% incidence and tumor susceptibility linked to multiple genetic loci, this model offers a relevant system for investigating HCC pathogenesis (16). In this study, we employed long-term HFD, high-iron (HID), and combined (HFD+HID) dietary interventions in C3HeB/FeJ mice to systematically characterize diet-specific and shared molecular alterations driving hepatocarcinogenesis.

Our findings reveal distinct and overlapping effects of HFD and HID on liver tumor progression, with HFD exerting a stronger influence on metabolic reprogramming and oxidative stress, while HID preferentially activates immune and inflammatory pathways. Transcriptomic profiling identifies conserved oncogenic programs and diet-specific signatures, including monotonic gene expression changes across disease stages that underscore the progressive nature of diet-induced liver carcinogenesis.

## Materials and methods

### Animals and dietary interventions

The wild-type inbred C3HeB/FeJ male and female mice were originally obtained from Jackson Laboratories and then bred in-house for the dietary intervention studies. All mouse experiments were approved by the Institutional Animal Care and Use Committee and monitored by the Department of Laboratory Animal Resources (DLAR) at the University of Texas Health Science Center at San Antonio. The mice were maintained in micro-isolator-topped cages and fed standard laboratory chow *ad libitum*. They were housed in pathogen-free facilities that are accredited by Association for the Assessment and Accreditation of Laboratory Animal Care (AAALAC). The health and behavior of animals were monitored daily by DLAR personnel. Two-month-old male C3HeB/FeJ mice fed *ad libitum* with a control diet (10 kcal% fat, 48 ppm iron, 5.7 ppm copper; TestDiet, Cat. # 1817636-203), HFD (61.1 kcal% fat, 48 ppm iron, 7.8 ppm copper; TestDiet, Cat. # 1817638-206), HID (10 kcal% fat, 200 ppm iron, 5.7 ppm copper; TestDiet, Cat. # 1817637-203), and HFD+HID (61.1 kcal% fat, 200 ppm iron, 7.8 ppm copper; TestDiet, Cat. # 1817639-206) for 16.5 months before termination. Details on fat and iron content of each diet are shown in Fig. S1a. At the termination, livers from euthanized mice were excised. Visible tumors on the surface of the livers were counted. Width (W) and length (L) of each tumor were measured with a ruler for calculation of its volume, which equals LxW^2^/2. Tumor containing liver tissues were carefully dissected and were either flash frozen or fixed in formalin along with their paired non-tumor liver tissues for further analysis.

### Quantification of iron in tissue and plasma samples

The iron content in liver tissues and plasma was measured by inductively coupled plasma mass spectrometry (ICP-MS, X Series II, Thermo Fisher, Waltham, MA). All samples (10-20 mg liver tissue or 100 µl plasma) were digested for 4□h in 1□mL of 70% nitric acid at 85°C. Samples were cooled at room temperature, centrifuged at 5,000 rpm for 1□min, diluted into 34□mL double-deionized water (2% nitric acid solution), vortexed, and assayed by ICP-MS as previously described (17).

### Western blotting

Total protein was extracted from xenograft tissues using Laemmli buffer with protease inhibitors. Western blotting was performed as previously described (18). Antibodies used in the study included anti-phospho-mTOR (Ser2448) (Cell Signaling, #2976), anti-mTOR (Cell Signaling, #2972), anti-FASN (Cell Signaling, #3180), anti-beta-actin (Invitrogen, #PIMA5 15739), anti-Mouse IgG (Jackson immunoresearch, #115-035-003) and anti-Rabbit IgG (Jackson immunoresearch, #111-035-003).

### RNA extraction and quantitative real-time PCR

Total RNA was extracted from liver tissues as previously described (18). One microgram of total RNA was reverse-transcribed (RT) to cDNA using random primers and M-MLV reverse transcriptase from Invitrogen Life Technology (Grand Island, NY). Quantitative real-time PCR was performed using SYBR Green PCR Mix from Life Technologies. All primers used in this study were synthesized by Integrated DNA Technologies (Coralville, IA). mTORC1 target mouse genes *Birc5* (Baculoviral IAP repeat-containing 5) and *Lpl* (lipoprotein lipase) were quantified by RT-PCR using *Birc5* forward primer: 5’-CGAGAACGAGCCTGATTTGG-3’, *Birc5* reverse primer: 5’-GCTCTCTGTCTGTCCAGTTTC-3’, *Lpl* forward primer: 5’-CATCTCATTCCTGGATTAGC-3’, and *Lpl* reverse primer: 5’-GTAGTAGACTGGTTGTATCGG-3’. Mouse *Gapdh* was used as an internal control and measured with forward primer: 5’-AGGTCGGTGTGAACGGATTTG-3’ and reverse primer: 5’-GGGCTGGTTGATGGCAACA-3’.

### Immunohistochemistry

Tissue was fixed for 24 hours in 10% neutral-buffered formalin, dehydrated in ethanol, and embedded in paraffin wax to make formalin fixed paraffin embedded (FFPE) tissue blocks. FFPE tissue sections were cut to 5 μm on glass slides, de-paraffined, and rehydrated by graded ethanol solutions. Antigen retrieval was executed by heating in sodium citrate (10 mM; pH 6.0; 95°C) for 10 minutes and allowed to return to room temperature for another 10 minutes. Endogenous peroxidase reaction was prevented by incubating sections with 3% hydrogen peroxide for 15 minutes. Nonspecific binding by antibodies was blocked with 10% goat serum for 30 minutes at room temperature. The sections were incubated with phospho-p70S6K1 (Thr389/412) (Invitrogen, #PA5-104842) overnight with phosphate-buffered saline and 0.025% Triton solution (PBST) and 5% goat serum in a humidified chamber at 4°C. Samples were then washed twice with PBST. Biotin-conjugated secondary antibody (Fisher Scientific, #BD550338) were incubated for 1 hour at room temperature. After washing twice with PBST, the samples were incubated with streptavidin-horseradish peroxidase for 30 min and counterstained with hematoxylin for 2 minutes before dehydration and mounting. Oxidized phospholipids (OxPLs) in FFPE tissue sections were stained using the E06 anti-OxPL antibody as previously described (19). IHC stained tissue was visualized with NanoZoomer Slide Scanner. OxPL staining was analyzed using ImageJ. Positive staining was quantified by thresholding pixel intensities. At least three randomly selected fields per sample were averaged for each mouse liver sample.

### RNA sequencing and bioinformatics analysis

To ensure fair comparisons, we selected an equal number of C3HeB/FeJ mice from each dietary group by randomly choosing four mice per group for RNA sequencing. Tumor and paired non-tumor liver samples from diet-induced HCC mice and normal liver samples from control diet-fed mice were used for genome-wide RNA sequencing.

Total RNA was extracted from the tumor, adjacent non-tumor, and normal liver samples obtained from male mice using the Qiagen RNeasy Mini Kit (Qiagen, Hilden, Germany) along with TRIzol reagent (Thermo Fisher Scientific, Waltham, MA). RNA sequencing libraries were prepared following the KAPA Stranded RNA-Seq Library Preparation protocol (KAPA Biosystems, Cape Town, South Africa). During the elution, fragmentation, and priming step, primer annealing was conducted at 85°C for 2 minutes to prevent RNA fragmentation. The quality of the extracted RNA was thoroughly evaluated using a Nanodrop spectrophotometer, 1% agarose gel electrophoresis, and a bioanalyzer. Poly-A mRNA was selectively enriched using magnetic beads conjugated with poly-T oligos. The enriched RNA was then fragmented and reverse-transcribed into first-strand cDNA using reverse transcriptase and random primers. Second-strand cDNA synthesis was carried out with DNA Polymerase I and RNase H. The resulting cDNA fragments underwent end repair, 3’-end adenylation, and adapter ligation. After purification, these libraries were amplified via PCR and sequenced as 50 bp single-end reads on the Illumina HiSeq 3000 platform at the UT Health San Antonio Genome Sequencing Core Facility.

The sequencing reads were aligned to the mouse genome (mm9) using TopHat2 (v2.0.8b)(20) with default parameters. Aligned BAM files were processed using the RSEM (v1.2.31)(21) algorithm to quantify gene expression levels, reporting expected counts and normalized expression values as reads per kilobase of transcript per million mapped reads (RPKM).

### Differential gene expression analysis and functional enrichment

Differentially expressed genes (DEGs) were identified from RSEM-derived counts using the DESeq2 package (v1.30.1)(22) in R/Bioconductor. Analyses focused on two comparisons: (a) paired tumor and adjacent non-tumor samples from C3HeB/FeJ mice subjected to high-fat, high-iron, or combined diets (denoted as TvNT), and (b) non-tumor samples compared to normal liver tissue from mice on a control diet (denoted as NTvN). For both comparisons, DEGs were identified as genes showing significant dysregulation (adjusted p-value < 0.05) with a minimum two-fold change (|log₂FC| ≥ 1) in tumor versus paired non-tumor tissue or in non-tumor versus normal liver samples.

Functional enrichment analysis for KEGG pathways and Gene Ontology (GO) Biological Process (BP) terms was conducted using the DAVID online platform(23). Enrichment analyses were performed separately for (a) all DEGs, (b) upregulated DEGs, and (c) downregulated DEGs. Pathways and GO BP terms with a false discovery rate (FDR) < 0.05 were considered significantly enriched.

### Gene set enrichment analysis (GSEA) and single-sample GSEA (ssGSEA)

The gene list for each HCC tumor sample from mice treated with (a) HFD, (b) HID, or (c) HFD+HID diets were ranked using the average log2 fold-change of expressed genes (average RPKM > 1) in tumors compared to paired non-tumor liver samples. Gene set enrichment analysis (GSEA) was performed with R package clusterProfiler (v3.18.1)(24) for assessing the enrichment of mouse MSigDB (v2024.1)(25) collections of hallmark gene sets, WikiPathways subset of canonical pathways, and Gene Transcription Regulation Database (GTRD)(26) gene sets. The GSEA analysis using clusterProfiler was conducted with all parameters set to their default configurations.

To explore the enrichment of hallmark gene sets (n=50) in each diet-induced HCC tumor sample we performed single-sample GSEA (ssGSEA) using the R-based ssGSEA2.0 package (https://github.com/broadinstitute/ssGSEA2.0).

### Statistical analysis

The effects of HFD), HID, and HFD+HID on body weight, tumor density, tumor volume in C3HeB/FeJ mice, and Ox-PL intensities were evaluated using one-way ANOVA. Paired t-tests were used to compare iron concentrations (ng/mg) between tumor and non-tumor tissues. For gene set enrichment analysis (GSEA), p-values for test statistics were calculated using permutation tests.

Partial least squares discriminant analysis (PLS-DA)(27) was applied to distinguish HCC tumors induced by different diets (HFD, HID, or HFD+HID). PLS-DA models were developed using the R Bioconductor package ropls (v. 1.22.0)(28), with pareto-scaled normalized enrichment scores (NES) from single-sample GSEA (ssGSEA) analyses of hallmark gene sets. Variable importance in projection (VIP) scores were used to evaluate the contribution of each gene set to the model, considering only gene sets with a VIP > 1 as potential biomarkers of diet-specific HCC induction in C3HeB mice.

To further analyze diet-specific effects, the Kruskal-Wallis test was used to assess overall significance across diets, and pairwise Wilcoxon tests were conducted to identify differences between any two diet groups. These combined statistical approaches provided a comprehensive analysis of the dietary impact on tumor development and molecular profiles.

## Results

### Impact of high-fat and high-iron diets on tumor development and iron distribution in C3HeB/FeJ mice

Two-month-old male C3HeB/FeJ mice were randomly assigned to one of four dietary regimens: a regular control diet (n=10), a HFD (n=9, after one death), a HID (n=7, after three deaths), or HFD+HID (n=10). The mice were maintained on these diets for 16.5 months (Fig. S1a). Throughout the treatment period, mice in the HFD, HID, and HFD+HID groups exhibited increased average body weight compared to controls (Fig. S1b). Notably, mice on the HFD alone or in combination with HID showed a statistically significant increase in body weight at the end of the treatment (p < 0.05, Fig. 1a), suggesting that the HFD was more effective than the HID in causing weight gain.

**Figure 1:**
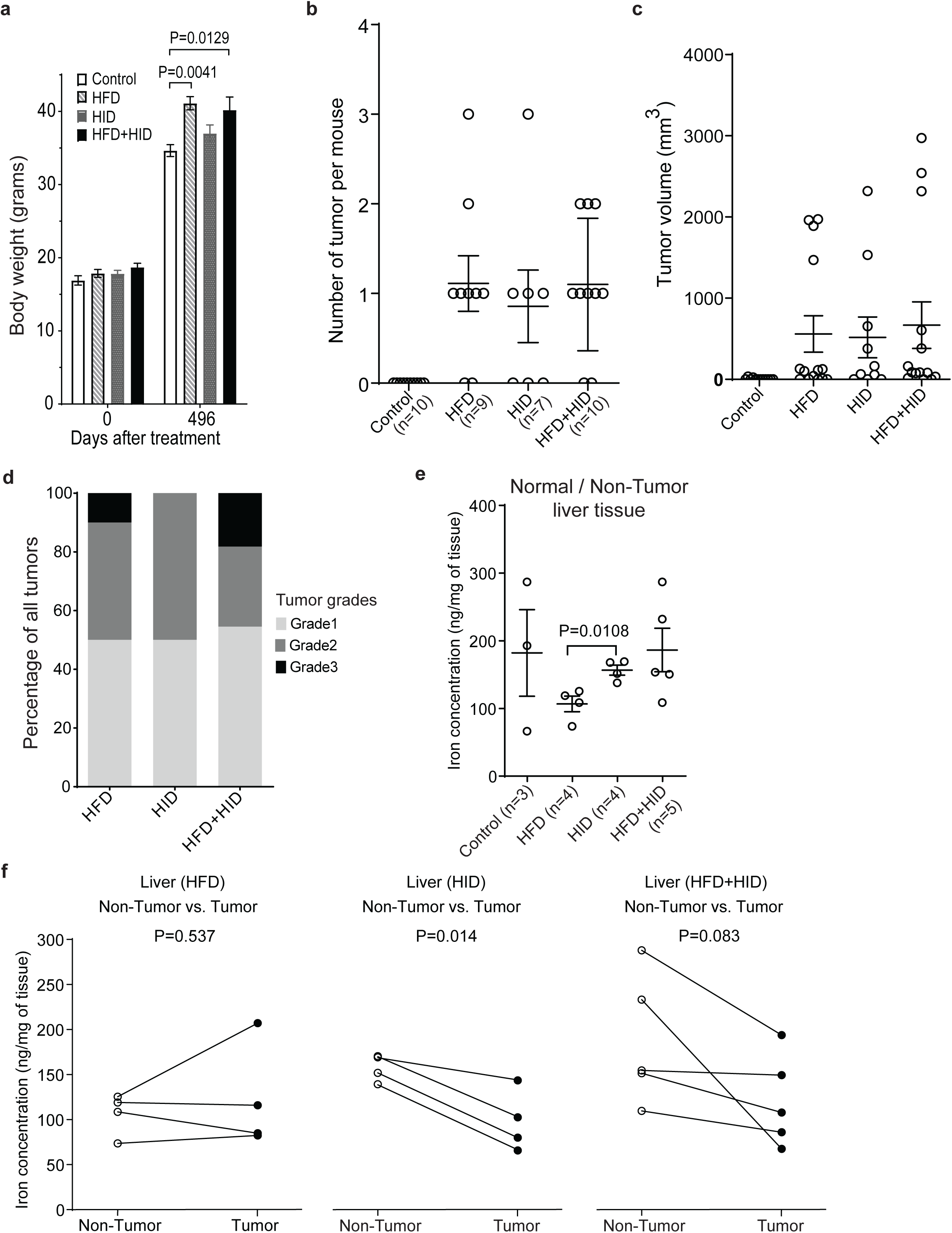
Effects of HFD and/or HID on body weight, tumor characteristics, and iron concentration in C3HeB/FeJ mice. (a) Mean body weight of mice treated with different diets. (b) Comparison of the number of liver tumors in individual mice across dietary groups. (c) Tumor volumes measured for all liver tumors (1–3 per mouse). (d) Tumor grade distribution. (e) Iron concentration (ng/mg tissue) in normal liver (control diet) compared to tumor-adjacent non-tumor liver samples from HFD, HID, and HFD+HID groups. (f) Paired comparison of iron concentration (ng/mg) between tumor and non-tumor tissues in HFD (left), HID (middle), and HFD+HID (right) groups. For (a-c, e): Data are expressed as mean ± SEM. Statistical analysis: For (a–c), one-way ANOVA; for (e–f), paired sample t-tests. Only significant p-values (p < 0.05) are shown. Abbreviations: HFD, high-fat diet; HID, high-iron diet; HFD+HID, combined high-fat and high-iron diets; SEM, standard error of the mean.

At the termination of the experiment, majority mice fed with HFD (77.8%), HID (57.1%), or HFD+HID (80%) developed HCC tumors (Fig. S1c) while the mice fed with the control diet had no tumor (Fig. 1b). There were no significant differences among the three diet groups in the number of tumors per mouse (Fig. 1b) or in tumor volume (Fig. 1c). Tumor grades, evaluated by Fisher’s exact test, were also comparable across the diet-induced HCC groups (Fig. 1d).

Iron concentration measurements in the normal liver from the mice fed with the control diet and the non-tumor liver tissues from the mice fed the three experimental diets revealed no significant differences among the four groups (Fig. 1e). The absence of statistical significance likely reflects substantial inter-sample variability in the control group together with limited statistical power, potentially obscuring diet-induced differences in hepatic iron levels. However, non-tumor samples from HID-fed mice showed significantly higher iron levels (p = 0.0108) than those from HFD-induced HCC (Fig. 1e). Plasma iron concentrations remained similar across all dietary groups even though most mice in the HFD+HID group had higher plasma iron levels (Fig. S1d). When comparing tumor tissues with their paired non-tumor samples, HFD-induced HCC showed comparable iron concentrations with no statistically significant differences (Fig. 1f, left). In contrast, tumors from HID-treated mice exhibited lower iron levels relative to adjacent non-tumor tissues, with a significant reduction observed for HID alone (p = 0.014) and a nearly significant reduction for the HID+HFD group (p = 0.083) (Fig. 1f, middle and right, respectively). This pattern suggests that the higher iron accumulated in the normal liver of the mice due to the high iron in their diet (Fig. 1e) was depleted in the tumors, likely for their increased metabolic demands, resulting in lower iron concentrations than the surrounding non-tumor liver tissue.

### Activation of mTOR signaling and accumulation of oxidized phospholipids in liver tumors

mTOR signaling pathway is known to be activated in cancers including HCC as a crucial regulator of cell growth, proliferation, and metabolism. Western blot analysis revealed a high level of phosphorylated mTOR in liver tumors in comparison to their paired non-tumor liver tissues (Fig. 2a), indicating a significantly elevated level of active mTOR signaling. IHC analysis of phosphorylated S6-kinase (p-S6K), a downstream target of mTORC1, showed no staining in normal liver tissues from control diet-treated mice (Fig. S2a), indicating an absence of proliferation under normal dietary conditions. In contrast, tumor tissues from all dietary treatment groups exhibited robust p-S6K immunoreactivity (Fig. S2b-d), confirming mTOR pathway activation in tumor cells. Paired non-tumor liver tissues showed intermediate staining levels, suggesting diet-induced alterations contributed to mTOR pathway modulation. FASN protein, a target of the mTOR signaling pathway, was increased in the tumors in comparison to their paired non-tumor tissues in the HFD-treated mice and to a lesser degree in HID-treated mice (Fig. 2a).

**Figure 2:**
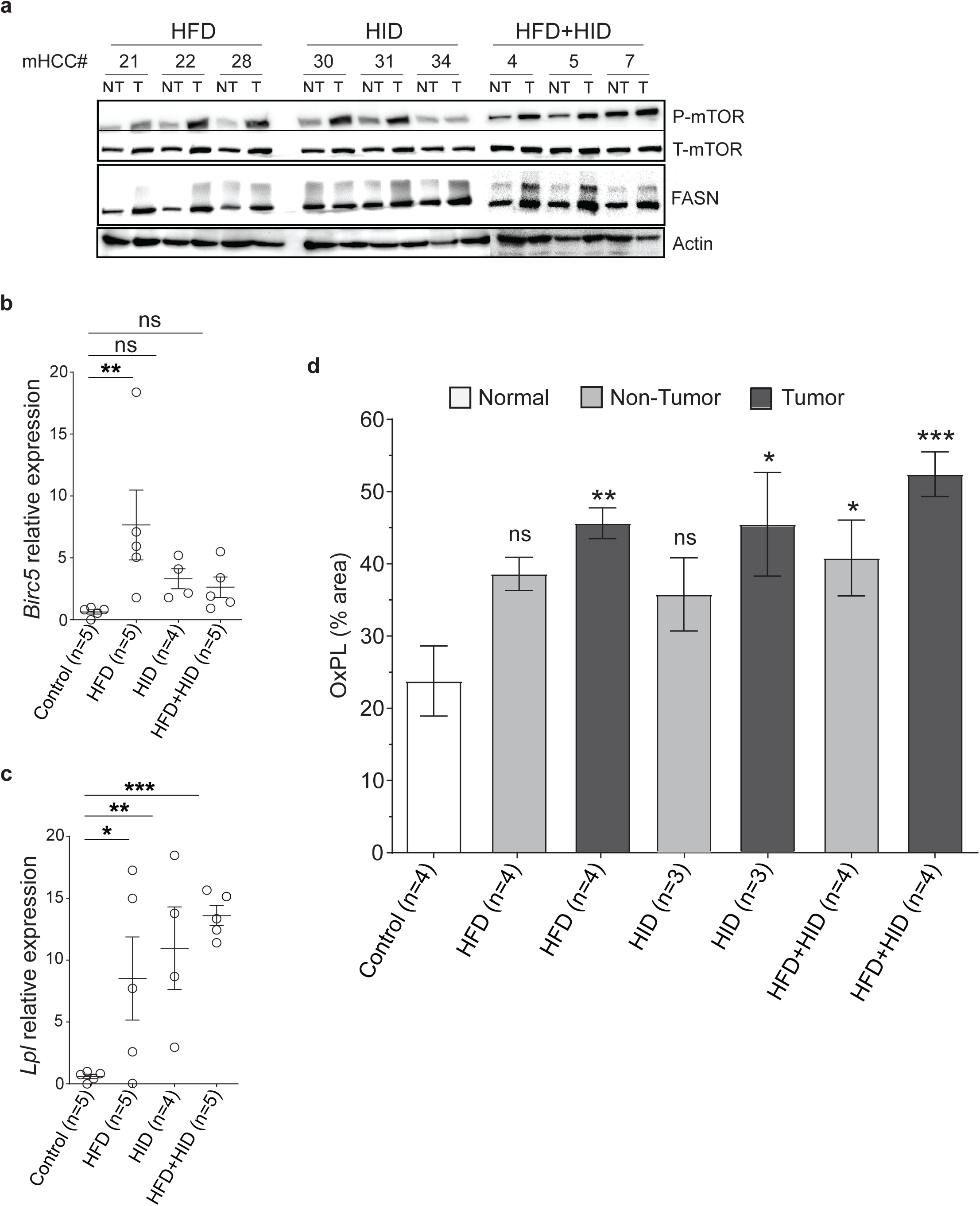
Effects of HFD and/or HID on mTOR signaling and oxidized phospholipid level. (a) Phospho-mTOR (P-mTOR), total mTOR (T-mTOR), and FASN protein levels in three paired non-tumor (NT) and tumor (T) samples from each diet treatment group were detected with Western blotting. The beta-actin protein levels were detected as total protein loading control. (b) Quantitative RT–PCR analysis of *Birc5* expression in control (normal) livers and tumors from HFD-, HID-, and HFD+HID-treated mice. Data are shown as relative expression normalized to *Gapdh*, with individual data points representing mice and bars indicating mean ± SEM. (c) Quantitative RT–PCR analysis of *Lpl* expression across the same groups as in (b), presented as relative expression (mean ± SEM). Statistical significance in (b) and (c) was assessed using ordinary one-way ANOVA comparing each treatment group with the control. (d) Liver sections were stained via immunohistochemistry (IHC) using an anti-OxPL antibody. The percent pixel intensity represents the level of positive staining. The plot shows the mean percent OxPL intensities ± SEM, calculated from the average of three different areas in each tissue section per mouse. OxPL intensities in normal liver (control diet) were compared to those in tumor and adjacent non-tumor liver samples from HFD, HID, and HFD+HID groups. Statistical significance was assessed using one-way ANOVA with Dunnett’s multiple comparison test. Significance levels for panels (b–d): ns = not significant, *P < 0.05, **P < 0.01, ***P<0.001.

We next quantified the expression of established mTOR-responsive genes at the transcript level. Quantitative real-time PCR analysis showed a significant induction of *Birc5* expression in tumors from HFD-treated mice with a moderate, not statistically significant, increase in HID-, and HFD+HID-treated mice compared with control livers (Fig. 2b). Notably, *Birc5* levels were highest in HFD-induced tumors, consistent with enhanced proliferative signaling under lipid-rich conditions. Similarly, expression of *Lpl*, a gene involved in lipid uptake and metabolism and regulated through mTOR-dependent metabolic programs, was significantly elevated in tumors across all diet-induced HCC models (Fig. 2c). These results indicate coordinated activation of mTOR-linked proliferative and metabolic gene expression in diet-induced liver tumors.

Oxidized phospholipids (OxPLs) accumulate under oxidative stress and are associated with MASH (19). Compared to normal liver tissues from control diet-treated mice, significantly elevated OxPL levels were observed in tumor tissues from all three diet treatment groups (Fig. 2d, S2e). Although OxPL accumulation in non-tumor tissues was higher than in normal liver tissues, statistical significance was only observed in the HFD+HID treatment group. These findings highlight unexpected similar effects of HFD and HID on mTOR pathway activation and oxidative stress as they induced liver tumorigenesis.

### Transcriptomic analysis unveils the profound influence of diet on liver tissue gene expression and tumorigenesis

Principal component analysis (PCA) of transcriptomic profiles (RPKM values) from liver tissue samples (Fig. 3a) demonstrated a clear separation between normal or adjacent non-tumor liver tissues and tumor samples from different diet-induced HCC models, indicating substantial transcriptomic shifts in HCC tumors. Notably, PCA of non-tumor liver tissues (Fig. 3b) revealed distinct clustering from normal liver tissues, highlighting diet-induced gene expression alterations even before tumorigenesis.

**Figure 3:**
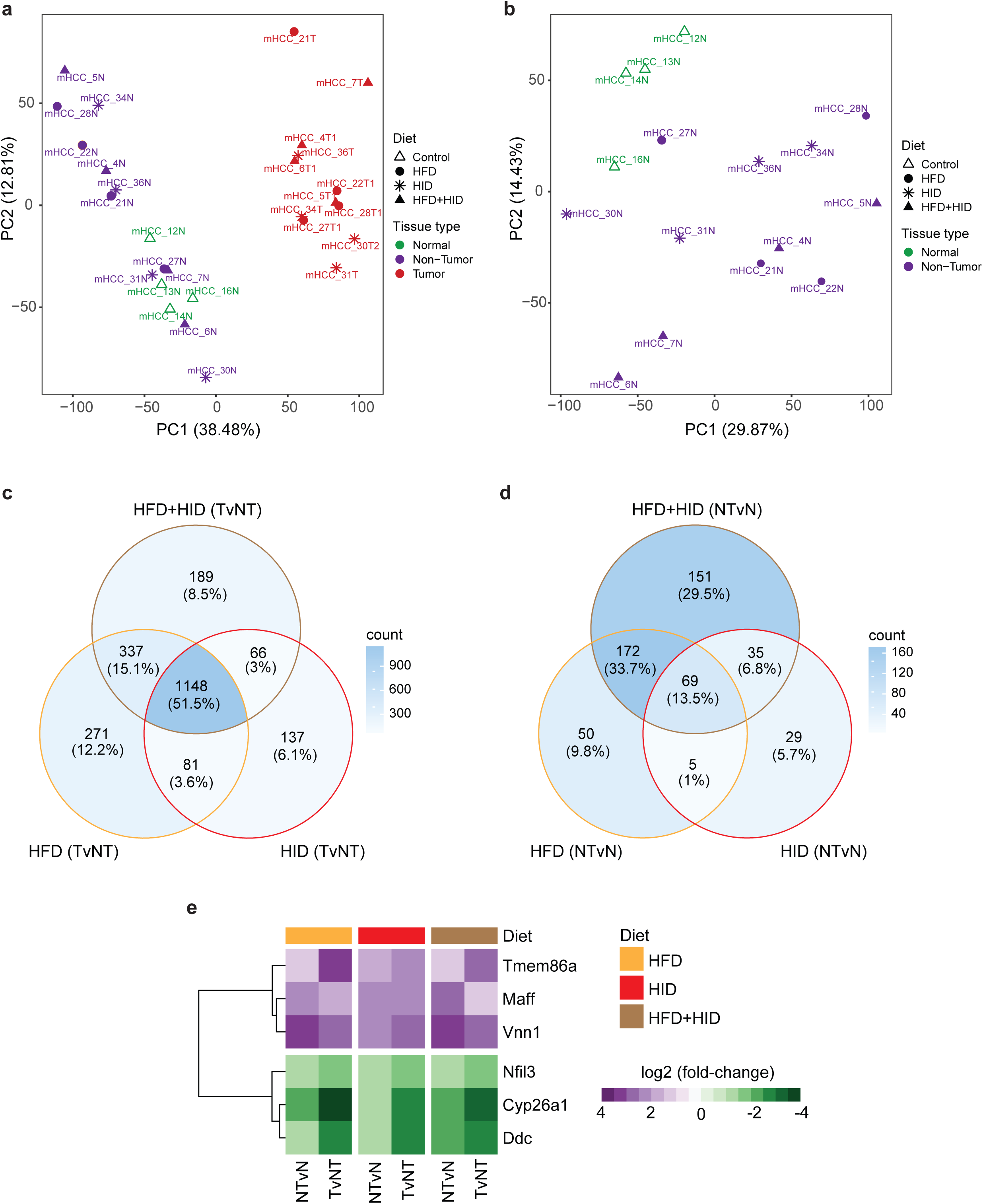
Transcriptomic analysis of liver tissues across dietary treatments. (a) Principal Component Analysis (PCA) plot of RNA sequencing data comparing normal liver tissue (control diet) and paired tumor and non-tumor tissues from HFD, HID, and HFD+HID-treated mice. (b) PCA plot comparing RNA sequencing data from normal liver tissues (control diet) to non-tumor liver tissues from HFD, HID, and HFD+HID groups. (c) Venn diagram illustrating the number of differentially expressed genes (DEGs) identified in tumor versus paired non-tumor tissue comparisons for HFD, HID, and HFD+HID-induced HCC. (d) Comparison of DEGs in non-tumor liver tissues from HFD, HID, and HFD+HID-treated mice versus normal liver tissues from the control diet group. (e) DEGs showing consistent monotonic upregulation or downregulation in non-tumor versus normal liver (NTvN) and tumor versus non-tumor tissue (TvNT) comparisons, regardless of dietary treatment.

A total of 1837, 1432, and 1740 significantly differentially expressed genes (DEGs) were identified in tumor versus adjacent non-tumor (TvNT) liver tissue comparisons from the HFD, HID, and HFD+HID models, respectively (Table S1–S3). Comparative analysis of these DEG sets revealed both shared and diet-specific transcriptomic alterations across the three diet-induced HCC models (Fig. 3c). Notably, approximately 52% of DEGs were commonly dysregulated across all models, with nearly equal distributions of upregulated and downregulated genes (54% vs. 49%) (Fig. S3a–b), suggesting a conserved transcriptomic response during liver tumorigenesis irrespective of dietary context.

In contrast, only 14% of DEGs overlapped between adjacent non-tumor and normal liver tissue (NTvN) comparisons (Table S4–S6; Fig. 3d), underscoring the tumor-specific nature of the observed transcriptional alterations. Notably, a greater proportion of DEGs were shared between the HFD and HFD+HID models (34%) than between the HID and HFD+HID models (7%), highlighting the dominant impact of high-fat intake on tumor transcriptomic profiles. The directionality of gene dysregulation was consistent with overall DEGs by HFD vs. HID, with comparable proportions of shared upregulated (32% vs. 6%) and downregulated (35% vs. 7%) genes (Fig. S3c–d).

While most genes exhibited unique up- or downregulation in tumor versus non-tumor comparisons, a subset overlapped between tumor vs. non-tumor and non-tumor vs. normal comparisons (Fig. S3e), highlighting shared molecular pathways disrupted by diet-induced liver damage and carcinogenesis. Notably, six DEGs—including *Tmem86a, Maff, Vnn1, Nfil3, Ddc*, and *Cyp26a1*—displayed a monotonic pattern of upregulation or downregulation across normal, adjacent non-tumor, and tumor tissues, independent of dietary conditions (Fig. 3e).

### Differential gene regulation across tumor and non-tumor liver tissues under dietary interventions

Among the 1148 DEGs (Fig. 3c) commonly dysregulated in tumors compared to non-tumor tissues across all three diet models, *Cyp3a44* exhibited a distinct dysregulation pattern in HID-treated HCC mice (Fig. 4a). While *Cyp3a44* was significantly upregulated in tumors of HFD- and HFD+HID-treated mice, it was downregulated in tumors compared to paired non-tumor samples in HID-treated mice, suggesting a diet-specific regulatory mechanism (Fig. 4a). Studies show that acetaminophen overdose, a leading cause of drug-induced acute liver failure, reduces the expression of several PXR (Pregnane X Receptor) target genes, including Cyp3a44, in the mouse liver(29). This suggests that HFD alone or HFD+HID may have a greater hepatotoxic impact than HID alone.

**Figure 4:**
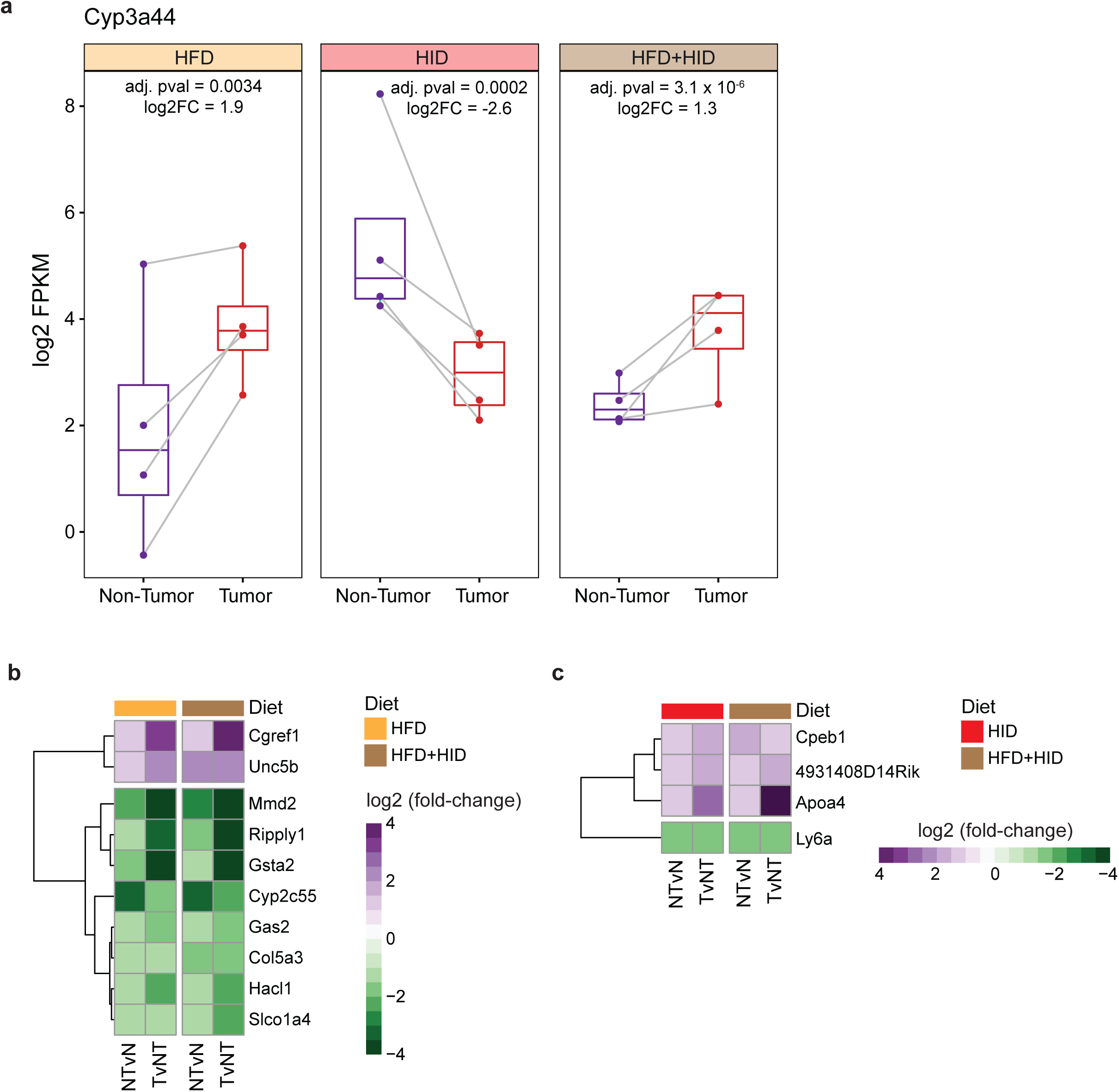
Differential regulation of common DEGs in tumor and non-tumor samples across dietary interventions. (a) Among the 1148 common differentially expressed genes (DEGs), *Cyp3a44* was the only gene showing opposite dysregulation in tumors compared to paired non-tumor samples with significant upregulation in HFD-or HFD+HID-treated mice, yet significant downregulation in HID-treated mice. (b) Common DEGs with significant and unidirectional dysregulation in non-tumor versus normal tissue (NTvN) and tumor versus non-tumor (TvNT) comparisons, demonstrating the potential effects of the high-fat diet on the liver. (c) A similar analysis for the high-iron diet identified DEGs significantly influenced by this dietary intervention.

Further analysis of common DEGs revealed subsets exhibiting significant and unidirectional dysregulation in tumor versus non-tumor (TvNT) and non-tumor versus normal tissue (NTvN) comparisons. These findings highlight the persistent impact of dietary factors on liver tissue remodeling, particularly under HFD conditions (Fig. 4b). A similar approach applied to HID-treated mice identified key DEGs uniquely influenced by high-iron dietary intervention, emphasizing its distinct molecular effects on liver pathophysiology (Fig. 4c).

Additionally, monotonic gene expression changes were observed across normal liver, non-tumor, and tumor tissues, reinforcing the progressive nature of gene dysregulation in diet-induced liver damage. A subset of DEGs showed consistent upregulation or downregulation in a diet-dependent manner, with unique signatures emerging in HFD-, HID-, and HFD+HID-treated mice (Fig. S4a–c). These findings suggest that specific dietary exposures drive distinct transcriptional alterations, potentially contributing to hepatocarcinogenesis through cumulative and pathway-specific effects.

### Enrichment of KEGG pathways and GO Biological Processes across dietary interventions

To understand the molecular mechanisms driving HCC under different dietary conditions, we performed pathway enrichment analyses using DEGs from tumor versus non-tumor comparisons across three dietary intervention models. This analysis revealed both shared and distinct enrichment patterns, offering insights into common and diet-specific pathways contributing to HCC progression.

Based on upregulated DEGs, 10 KEGG (Kyoto Encyclopedia of Genes and Genomes) pathways were consistently enriched across all three dietary groups (Fig. 5a, e), reflecting activation of conserved oncogenic signaling regardless of dietary background. Notable pathways included ECM-receptor interaction, pathways in cancer, PI3K-Akt signaling, and cytokine-cytokine receptor interaction. Complementary GO Biological Process (GOBP) analysis (Fig. 5b, f, and S5a) revealed enrichment of tumor-promoting biological processes such as lipid metabolism—notably the lipid catabolic process (GO:0016042 in Fig. S5a), angiogenesis, positive regulation of cell migration, apoptotic process, and cell adhesion. These findings suggest that across dietary conditions, upregulated DEGs converge on common biological programs that promote proliferation, angiogenesis, migration, and metabolic rewiring in HCC.

**Figure 5:**
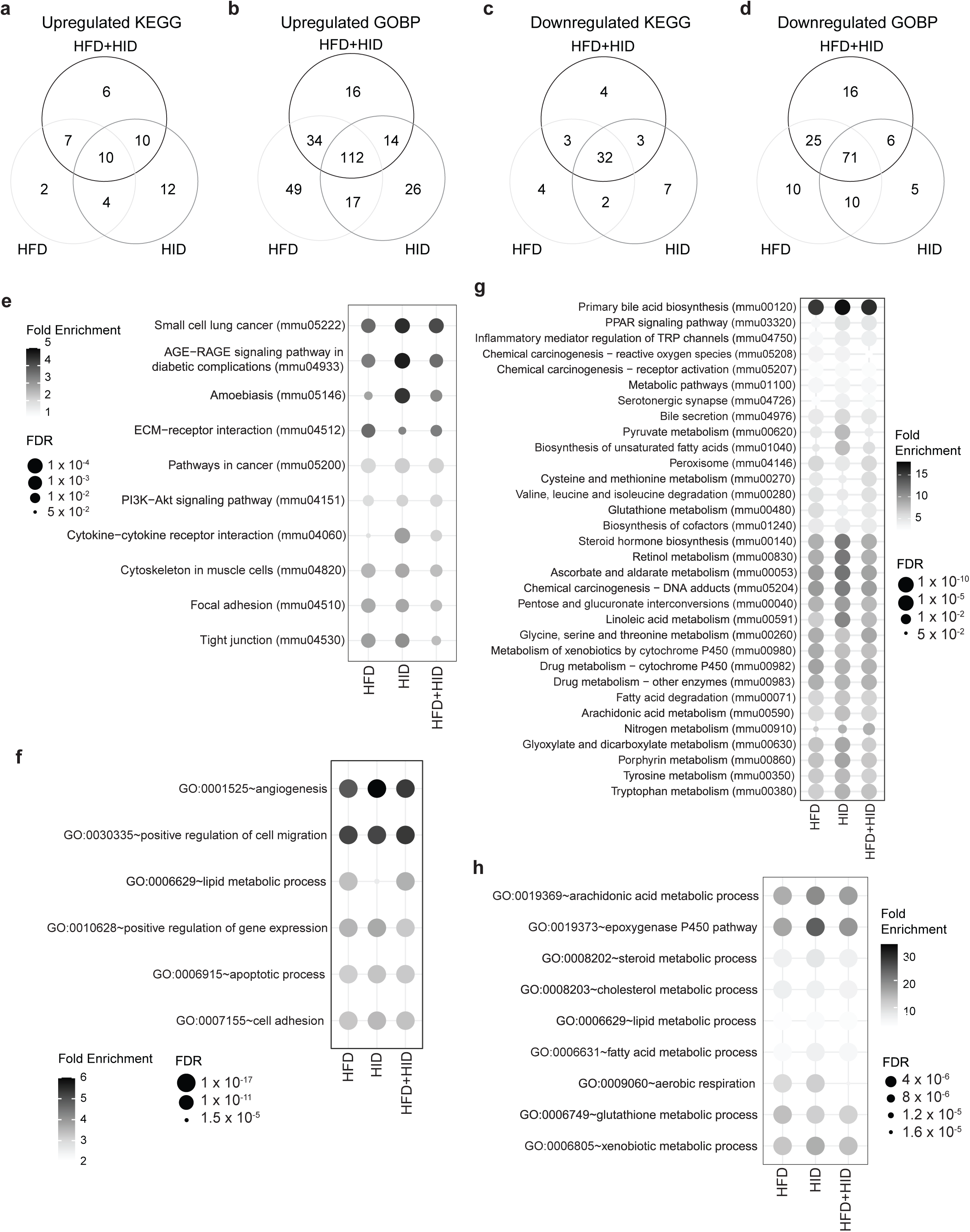
Comprehensive enrichment analysis of KEGG pathways and GO Biological Processes for differentially expressed genes across dietary interventions. (a, b) Venn diagrams illustrating the overlap of significantly enriched (a) KEGG pathways and (b) GO Biological Processes (GOBP) due to upregulated DEGs in tumor versus non-tumor comparisons across HFD, HID, and HFD+HID-induced HCC models. (c, d) Venn diagrams showing significantly enriched (c) KEGG pathways and (d) GOBP associated with downregulated DEGs across the three dietary groups. (e) Ten KEGG pathways commonly enriched with a false discovery rate (FDR) less than 0.05 by upregulated DEGs. (f) Six most significantly enriched GO Biological Processes associated with upregulated DEGs. (g) Thirty-two KEGG pathways commonly enriched by downregulated DEGs. (h) Nine most significantly enriched GO Biological Processes associated with downregulated DEGs. Abbreviations: KEGG, Kyoto Encyclopedia of Genes and Genomes; GO, Gene Ontology; GOBP, GO Biological Processes.

Conversely, downregulated DEGs showed consistent suppression of 32 KEGG pathways across all dietary models (Fig. 5c, g), including pathways associated with PPAR signaling, xenobiotic metabolism, retinol metabolism, and other liver-specific functions. GOBP enrichment (Fig. 5d, h, and S5b) highlighted significant downregulation of processes critical for hepatic homeostasis, such as aerobic respiration, arachidonic acid metabolic process, xenobiotic catabolic process (GO:0042178), and positive regulation of lipid metabolic process (GO:0045834 in Fig. S5a). This reflects a disruption of hepatocyte metabolic integrity and detoxification capacity during tumorigenesis across the different diet conditions.

### Enrichment of Distinct Pathways and Biological Process by Different Diets in HCC Development

Among the pathways significantly enriched (FDR < 0.05 or -log_10_ FDR > 1.301) due to upregulated DEGs in tumors revealed unique KEGG pathways (Fig. 6a) and GOBP terms (Fig. S6a, b, c) enriched for each dietary intervention. The HID-only tumors exhibited significant enrichment of several immune-related pathways including TNF (mmu04668) / IL-17 (mmu04657) / Toll-like receptor (mmu04620) signaling pathways (Fig. 6a), positive regulation of T cell activation (GO:0050870) / proliferation (GO:0042102) (Fig. S6b). Ras signaling pathway (mmu04014) was found significantly enriched in combination diets (Fig. 6a).

**Figure 6:**
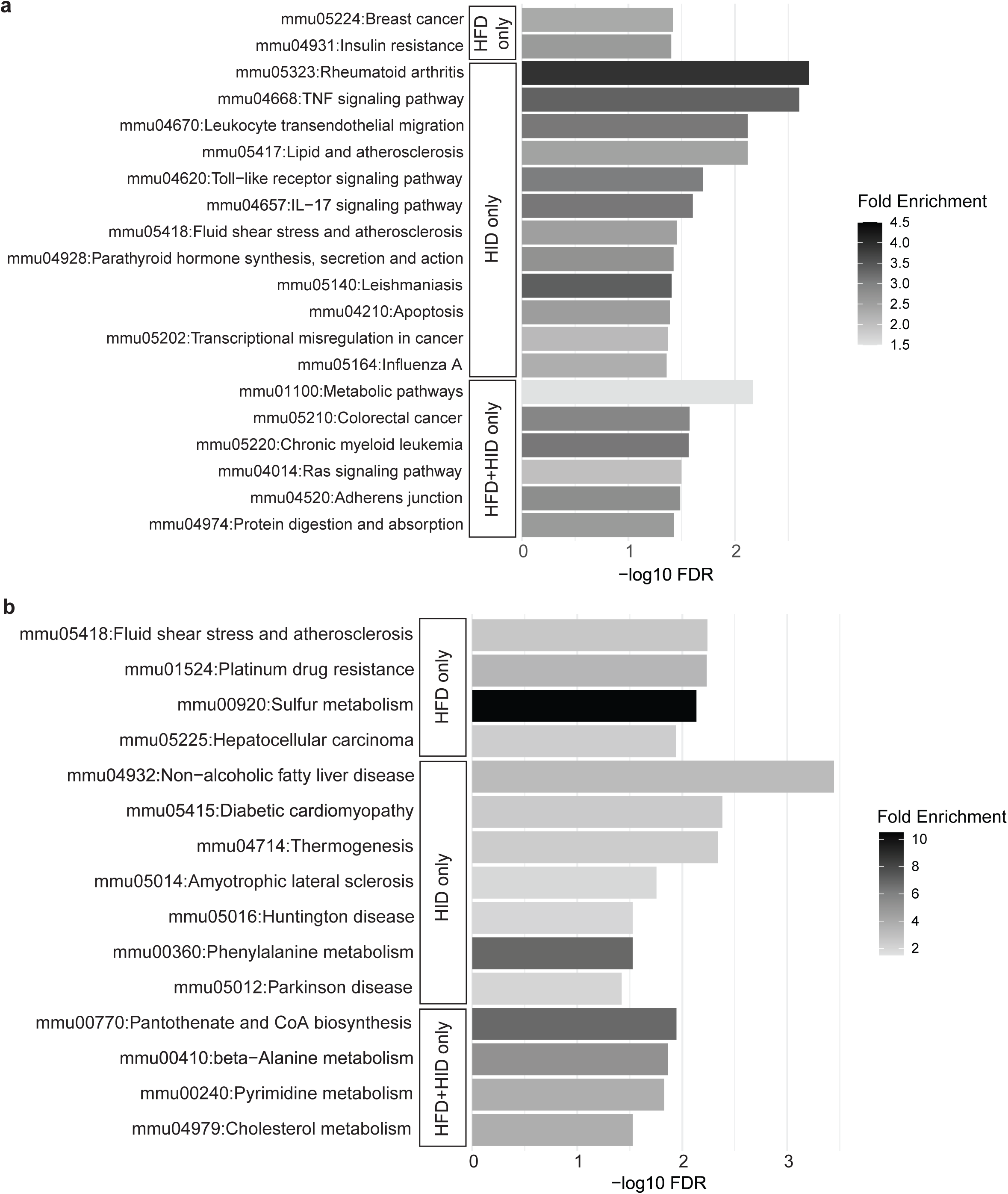
KEGG pathway enrichment analysis of differentially expressed genes due to specific dietary interventions on HCC development. (a) KEGG pathways significantly enriched due to upregulated DEGs in tumors compared to non-tumor samples, specific to each diet: HFD-only (n=2), HID-only (n=12), and HFD+HID-only (n=6). (b) KEGG pathways significantly enriched due to downregulated DEGs in tumors compared to non-tumor samples, specific to each diet: HFD-only (n=4), HID-only (n=7), and HFD+HID-only (n=4). These findings emphasize the unique molecular effects of each dietary regimen on the development of hepatocellular carcinoma (HCC).

Significantly enriched pathways using downregulated DEGs revealed enrichment of different metabolism related pathways uniquely regulated by any one diet or combination diets (Fig. 6b, S6d, e, f). In HFD diet alone, sulfur metabolism (mmu00920) as well as hydrogen sulfide biosynthetic process (GO:0070814) were found significantly enriched (Fig. 6b, S6d). Sulfur metabolism integrates amino acid catabolism and methylation cycles with hydrogen sulfide (H2S) synthesis, underscoring its role in redox balance and cellular signaling(30).

These results point to both shared and distinct biological processes perturbed in response to specific dietary treatments, offering insight into the mechanistic similarities and heterogeneity of diet-induced HCC. Together, these enrichment profiles demonstrate that the type and combination of dietary supplementations distinctly shape the transcriptomic landscape of HCC, influencing immune, metabolic and oncogenic signaling pathways.

### Diet-Specific Gene Set Enrichment Reveals Distinct Immune and Metabolic Signatures in HCC

Gene set enrichment analyses using hallmark and WikiPathways (WP) gene sets from MSigDB revealed both shared oncogenic programs and diet-specific transcriptomic alterations in tumor tissues compared to paired non-tumor controls (Fig. 7a, S7a). Metabolic gene sets, including hallmark bile acid metabolism, fatty acid metabolism, and WP amino acid metabolism (notably tryptophan metabolism), were consistently downregulated across all diet-induced HCCs. In contrast, hallmark adipogenesis and the WP TCA cycle were uniquely suppressed in HID-induced tumors. At the same time, the WP oxidative stress and redox pathway was significantly downregulated in HFD and HFD+HID groups (Fig. 7a, S7a).

**Figure 7:**
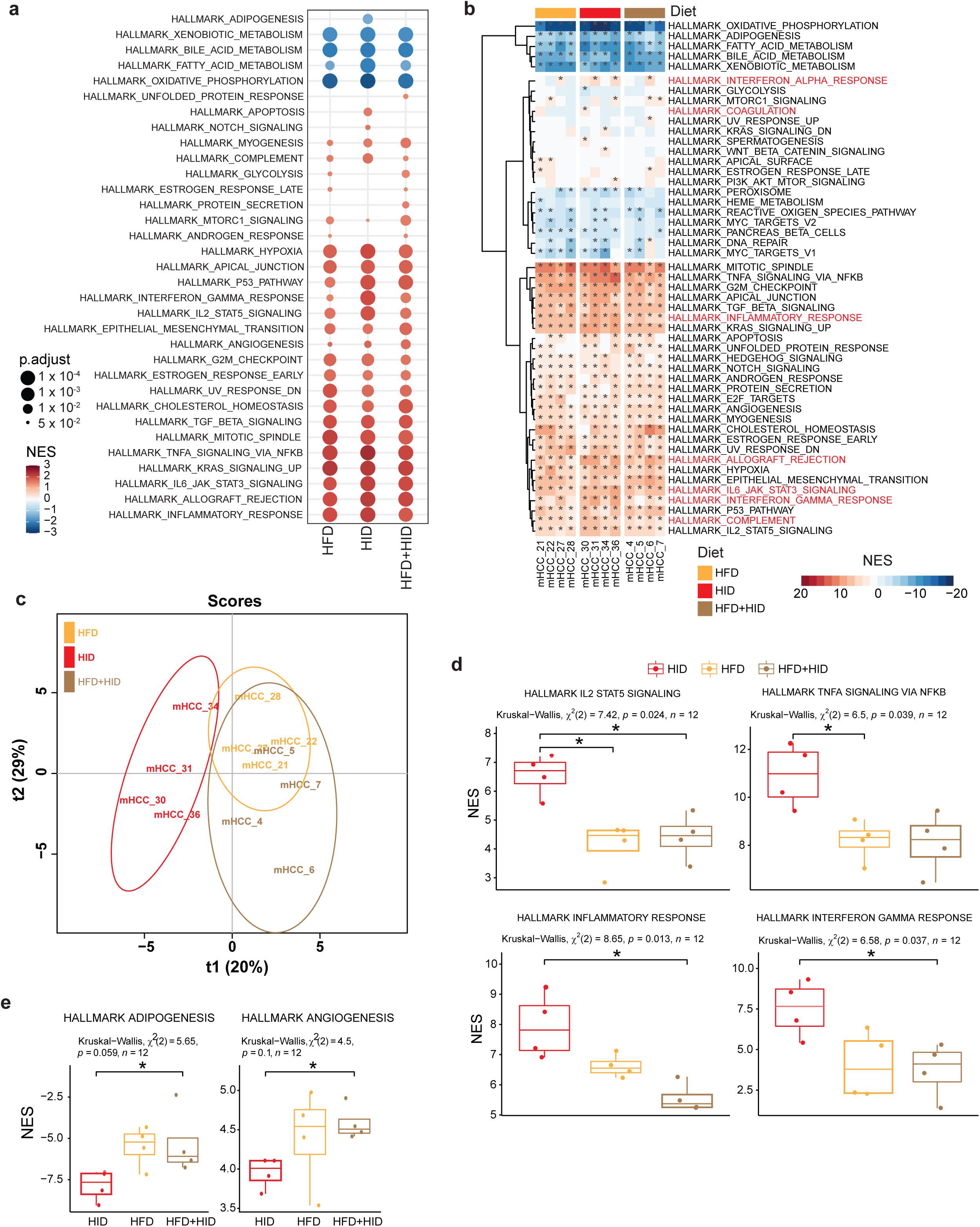
Similar and distinct hallmark gene set enrichments due to dietary interventions on HCC development. (a) Hallmark gene sets significantly enriched in tumors compared to non-tumor samples specific to HFD, HID, and HFD+HID. (b) Enriched hallmark gene sets in individual tumor samples relative to paired non-tumor samples in each diet group. Asterisks denote significant enrichment (FDR < 0.05), and seven immune-related hallmark gene sets are highlighted in red. (c) Partial least squares discriminant analysis (PLS-DA) score plot based on normalized enrichment scores (NES) of the 50 hallmark gene sets, reflecting different effects of diets on gene set enrichments in tumors. (d) Four gene sets with significantly different enrichments across dietary groups (Kruskal-Wallis test, p < 0.05) showed consistently higher enrichments in HID-induced HCC compared to HFD or HFD+HID diet-induced HCC. (e) Hallmark adipogenesis and angiogenesis gene sets exhibited significantly lower enrichment in HID-induced HCC compared to HFD+HID-induced HCC. Statistical significance in (d) and (e) was determined using the Wilcoxon test (*P < 0.05).

Proliferative signatures, such as hallmark cell cycle-related gene sets, were significantly upregulated across all diets, reflecting sustained cell cycle activity in tumor tissues. Likewise, immune-related hallmark gene sets were broadly upregulated, indicating an immune-infiltrated tumor microenvironment (Fig. 7a). Notably, WP IL-5 signaling and type II interferon signaling (IFNG) showed more substantial enrichment in HID-induced tumors (Fig. S7a), suggesting heightened immune activation under iron overload. Supporting this, transcriptional targets of Rel/NF-κB, a key immune regulator (31,32), were significantly enriched in HID tumors (Fig. S7b), pointing to HID-specific activation of NF–κB–mediated immune pathways.

Individual sample-level analysis of hallmark gene sets revealed generally consistent enrichment patterns with modest interindividual variation (Fig. 7b). Except for coagulation, interferon-α response, and interferon-γ response, four immune-related hallmark gene sets were significantly positively enriched (FDR < 0.05, NES > 0) in all 12 mouse tumor samples relative to their paired non-tumor liver tissues (Fig. 7b). Cross-species clustering of tumors from 13 Hispanic HCC patients(33) and 12 diet-induced mouse tumors revealed that C3HeB/FeJ mice clustered with the immunologically “hot” human HCC (HM-1) subgroup (Fig. S7c), reinforcing the translational relevance of this model for studying immune-enriched tumor phenotypes.

To investigate intergroup differences, PLS-DA of normalized enrichment scores across 50 hallmark gene sets revealed clear separation among HFD, HID, and HFD+HID tumor profiles (Fig. 7c, Table S7), confirming diet-specific transcriptomic signatures. Notably, inflammatory response and interferon signaling gene sets showed significantly higher enrichment (*p* < 0.05, Kruskal-Wallis test) in HID-induced tumors (Fig. 7d), consistent with enhanced immune activation. In contrast, adipogenesis and angiogenesis were significantly less enriched in HID tumors compared to HFD+HID tumors (Fig. 7e, *p* < 0.05, Wilcoxon test), suggesting suppression of lipid and vascular remodeling pathways in iron-driven hepatocarcinogenesis.

Together, these results demonstrate that while core oncogenic pathways are commonly activated in diet-induced HCC, distinct immune and metabolic responses are modulated by specific dietary exposures. Among them, HID induces the most pronounced immune activation signaling, highlighting its higher impact on shaping the tumor immune microenvironment.

## Discussion

Our present study demonstrates that long-term consumption of HFD and/or HID promotes HCC through divergent yet partially overlapping molecular programs, involving iron metabolism, mTOR signaling, oxidative stress, and transcriptomic reprogramming

Consistent with previous reports, HFD with or without HID significantly increased body weight, yet tumor number and volume were comparable across all diet groups, indicating that HFD promotes metabolic dysregulation without disproportionately amplifying overall tumor burden in this model.

Male C3HeB/FeJ mice were known to develop spontaneous HCC (15,16). However, the mice fed with the control diet did not develop HCC in our study. The previous observations are likely due to high iron levels in standard laboratory mouse chow used in earlier studies. For example, the Teklad LM-485 mouse diet (Harlan Laboratories, Cat. # 7012) used by Zavadil and co-workers (16) contained 240 ppm iron, similar to that in the HID used in our study. Accordingly, the HID mice in the present study were fed a diet containing 200 ppm iron, a level similar to that of standard commercial rodent chow and substantially higher than many defined low-iron purified diets, thereby supporting its classification as an iron-enriched dietary condition (34).

Iron quantification revealed diet- and tissue-specific changes in hepatic iron. Non-tumor tissues from HID-fed mice had significantly higher iron than those from HFD-fed mice, consistent with dietary iron loading; plasma iron was unchanged across groups. Strikingly, tumors from HID-fed mice showed significantly lower iron than paired non-tumor tissue, while HFD tumors maintained comparable levels. This intratumoral iron depletion suggests active iron utilization or redistribution to meet the heightened metabolic demands of HID-driven tumors. In a liver-specific Trp53 deficient mouse model of with CCl₄-induced liver carcinogenesis, HCC tumors also exhibited significantly lower iron concentrations compared to adjacent non-tumor liver tissues in both male and female mice (35). In addition, livers of HFD-induced HCC in mice with and without maternal exposure to cadmium also showed low levels of iron compared to control mice without HFD treatment (17). One potential assumption is that ferroptosis, an iron-dependent lipid peroxidation-mediated cell death, in the liver possesses a significant anticancer role by acting as a form of regulated cell death that eliminates HCC cells (36,37). Due to the chronic decrease iron in the liver, the reduced capacity of anticancer ferroptotic removal of the developed cancer cells may be a potential link. However, it should be noted that the observed reduction in tumor iron levels may not reflect solely active consumption, but could also indicate redistribution to other compartments or altered iron uptake and storage dynamics within the tumor microenvironment; therefore, the interpretation of tumor iron depletion should be considered with caution.

The activation of mTOR signaling in all tumor tissues, as evidenced by increased phosphorylation of S6K and mTOR, underscores a conserved mechanism of oncogenic signaling in diet-induced HCC (38). Since mTOR integrates growth signals with nutrient availability, its activation likely reflects the nutrient-rich environment created by high-fat or iron-rich diets. Concordantly, the upregulation of *Birc5* and *Lpl* across diet-induced tumors supports coordinated activation of mTOR-driven proliferative and metabolic programs. Elevated *Birc5*, particularly in HFD-induced tumors, is consistent with enhanced proliferative signaling under lipid-rich conditions (39), while increased *Lpl* expression suggests augmented lipid uptake and metabolic flexibility in tumor cells (40). Elevated OxPL levels in HFD and HFD+HID tumors further implicate dietary fat in driving lipid peroxidation and oxidative stress, consistent with impaired redox balance in these tumors (41). OxPL accumulation in pre-neoplastic non-tumor tissues further suggests that lipid oxidative stress precedes overt tumorigenesis (42).

Transcriptomic analysis revealed extensive gene expression changes across dietary groups, with a clear separation between normal, non-tumor, and tumor tissues. Remarkably, even non-tumor tissues clustered separately from normal livers, suggesting that dietary exposure induces substantial gene regulatory shifts before tumor formation. Shared differentially expressed genes (DEGs) among tumor groups point to conserved transcriptional programs involved in cell cycle regulation, immune modulation, and metabolic reprogramming. Notably, the substantially greater overlap of DEGs between HFD and HFD+HID groups (34%) compared to HID and HFD+HID (7%) underscores the dominant transcriptional influence of dietary fat.

A subset of DEGs exhibited monotonic expression changes across normal, adjacent non-tumor, and tumor tissues irrespective of diet, suggesting their role in progressive molecular remodeling during hepatocarcinogenesis. Notably, Tmem86a, which functions as a lysoplasmalogenase and LXR target, has been implicated in lipid metabolism and membrane remodeling under high-fat diet–induced stress (43,44). MAFF, enriched in liver tumors and tumor-initiating cells, is a proposed therapeutic target, with antisense oligonucleotides demonstrating anti-cancer efficacy (45). VNN1, a liver-enriched pantetheinase, regulates gluconeogenesis and is upregulated in obese mouse models (46). On the other hand, the downregulation of Nfil3 was shown to enhances hepatic gluconeogenic gene expression, supporting glucose supply for proliferating tumor cells (47). DDC, downregulated in laryngeal cancer (48), may similarly mark tumor progression in liver. Cyp26a1 downregulation by metformin was shown to increase retinoic acid levels and attenuate hepatocarcinogenesis in Fah⁻/⁻ mice (49) and therefore, it is not clear why it was downregulated during HFD- and/or HID-induced hepatocarcinogenesis. Collectively, these genes represent potential biomarkers and therapeutic targets involved in early-to-late stages of liver disease progression.

Our transcriptomic analysis reveals 1,148 DEGs shared across all three diet-induced HCC models. Among these, *Cyp3a44*, a gene regulated by the pregnane X receptor (PXR) and involved in xenobiotic metabolism (50), exhibited opposing expression patterns under HID and HFD conditions. Downregulation of *Cyp3a44* in HID-treated mice, contrasted with upregulation under HFD, suggests that high-fat diets may elicit greater hepatotoxic stress than iron overload, likely driven by enhanced lipid peroxidation and metabolic burden (51,52).

Monotonic gene expression trends observed across normal, non-tumor, and tumor tissue suggest sustained transcriptional dysregulation during the progression of hepatocellular carcinoma. In both the HFD and HFD+HID groups, genes such as *Cgref1* and *Unc5b* were progressively upregulated, whereas *Mmd2*, *Ripply1*, *Gsta2*, *Cyp2c55*, *Gas2*, *Col5a3*, *Hacl1*, and *Slco1a4* were consistently downregulated. Notably, *CGREF1* has been reported to promote HCC cell proliferation, invasion, and migration through the upregulation of *EIF3H* and subsequent activation of the Wnt/β-catenin signaling pathway (53). Similarly, *UNC5B*, a netrin-1 receptor, although extensively studied in breast cancer, is known to be associated with tumor progression and poor survival outcomes when overexpressed (54). Among the downregulated genes, *GAS2* (Growth arrest-specific 2) has been implicated in the suppression of HCC cell proliferation; its knockdown facilitates hepatocyte proliferation, emphasizing its role as a potential tumor suppressor in hepatic tissue (55).

In contrast, the HID group exhibited a distinct transcriptional signature characterized by elevated expression of *Cpeb1*, *Apoa4*, and *4931408D14Rik*, along with downregulation of *Ly6a*. These alterations likely reflect iron-induced stress impacting hepatic regeneration, redox homeostasis, and post-transcriptional control. *CPEB1* (cytoplasmic polyadenylation element-binding protein 1), a post-transcriptional regulatory factor, is implicated in enhancing erastin-induced ferroptosis (56). *APOA4*, an apolipoprotein involved in lipid metabolism and anti-inflammatory responses, has also been shown to be responsive to dietary and oxidative stress cues in hepatic tissue (57). Downregulation of *Ly6a* (also known as Sca-1), a stemness and immune-modulatory marker, suggests altered regenerative signaling and stem/progenitor cell dynamics under iron overload conditions (58). These findings reveal distinct, diet-induced molecular trajectories of HCC progression and implicate specific gene regulatory networks linked to lipid and iron metabolism, detoxification, and extracellular matrix (ECM) remodeling.

Integrated KEGG and GO pathway enrichment analyses revealed both shared and diet-specific molecular mechanisms underlying HCC progression. Across all dietary groups, tumors consistently exhibited upregulation of core oncogenic pathways, including PI3K–Akt signaling, ECM–receptor interaction, and lipid catabolism, indicating a conserved molecular foundation for tumor development irrespective of dietary context (59,60). In parallel, pathways critical for hepatic detoxification and mitochondrial function—such as xenobiotic metabolism, peroxisome proliferator-activated receptor (PPAR) signaling, and oxidative phosphorylation—were uniformly downregulated, highlighting a profound disruption of normal liver homeostasis during tumorigenesis (61,62). Consistent with these observations, gene set enrichment analysis (GSEA) revealed shared transcriptomic reprogramming across dietary models, characterized by coordinated suppression of bile acid, fatty acid, and tryptophan metabolism (63). These metabolic alterations represent established hallmarks of HCC, with impaired bile acid metabolism linked to poor prognosis (64) and dysregulated fatty acid and tryptophan metabolism contributing to tumor growth, immune modulation, and metabolic adaptation (65). Collectively, these conserved oncogenic and metabolic signatures underscore fundamental mechanisms of liver cancer development and suggest therapeutic vulnerabilities that transcend dietary etiology.

Despite this shared oncogenic core, distinct diet-specific pathway signatures were also evident. Tumors arising under an iron-rich diet (HID) showed selective enrichment of immune and inflammatory signaling pathways, including TNF, IL-17, and NF-κB signaling, as well as hallmark gene sets associated with apoptosis (66,67). These findings suggest that iron overload promotes a pro-inflammatory tumor microenvironment and may trigger apoptotic programs as a compensatory response to iron-induced oxidative stress. HID-induced tumors also displayed unique suppression of adipogenesis (68) and the tricarboxylic acid (TCA) cycle, consistent with mitochondrial and bioenergetic rewiring, which is a recognized metabolic feature of HCC that supports tumor survival and biosynthetic demands (69). In contrast, tumors from high-fat diet (HFD) and combined HFD+HID models exhibited stronger enrichment of pathways related to oxidative stress responses, unfolded protein response, and lipid metabolic reprogramming, reflecting metabolic and endoplasmic reticulum stress associated with nutrient excess (70). Notably, oxidative stress pathways were significantly downregulated in these tumors, a finding consistent with adaptive redox remodeling observed in obesity-associated HCC, in which dysregulated reactive oxygen species signaling contributes to tumor progression (71). Together, these results highlight how distinct dietary pressures shape unique immune and metabolic adaptations superimposed on a shared oncogenic framework in HCC.

Importantly, cross-species clustering analysis revealed that tumors from C3HeB/FeJ mice aligned with immunologically “hot” human HCC subtypes, supporting the translational relevance of this model (33,72).

Our findings illustrate that while high-fat and high-iron diets both contribute to liver tumorigenesis, they do so via distinct molecular routes. High-fat diets predominantly promote metabolic reprogramming, lipid oxidation, and mTOR activation, while high-iron diets trigger immune activation and pro-inflammatory rewiring. Their combination generates a transcriptomic landscape not reducible to either diet alone, with implications for prevention and therapeutic targeting in metabolic and iron-associated HCC.

## Data Availability

The RNA sequencing (RNA-seq) data generated in this study are being deposited in the Gene Expression Omnibus (GEO) repository. The accession number will be provided upon completion of the submission and will be made publicly available prior to publication. All other data supporting the findings of this study are available within the article and its Supplementary Information files. Additional data and materials are available from the corresponding author upon reasonable request.

## Author Contributions

Conceptualization: DD, HB, SPA, FGC, L-ZS; Funding acquisition and project administration: FGC, L-ZS; Animal experiments: HB; Investigation: all authors; Pathology: FES; Methodology: DD, HB, XS, JX, LC, L-ZS; Data analysis: DD, HB, XS, JX, LC, YC, L-ZS; Writing—original draft: DD, HB, L-ZS; Writing—review and editing: all authors.

## Acknowledgements

We thank Mr. Kyle Pressley and Mr. Kalyan Kakarla for their assistance in feeding the mice with the various diets and collecting mouse tissues.

## References

1. Bray F, Laversanne M, Sung H, Ferlay J, Siegel RL, Soerjomataram I, et al. Global cancer statistics 2022: GLOBOCAN estimates of incidence and mortality worldwide for 36 cancers in 185 countries. CA Cancer J. Clin. 2024;74:229–263.

2. McGlynn KA, Petrick JL, El-Serag HB. Epidemiology of Hepatocellular Carcinoma. Hepatology. 2021;73 Suppl 1:4–13.

3. Tian Y, Wong VW-S, Chan HL-Y, Cheng AS-L. Epigenetic regulation of hepatocellular carcinoma in non-alcoholic fatty liver disease. Semin. Cancer Biol. 2013;23:471–82.

4. Kowdley K V. Iron, hemochromatosis, and hepatocellular carcinoma. Gastroenterology. 2004;127:S79–S86.

5. Gan C, Yuan Y, Shen H, Gao J, Kong X, Che Z, et al. Liver diseases: epidemiology, causes, trends and predictions. Signal Transduct. Target. Ther. 2025;10:33.

6. Tsuru H, Osaka M, Hiraoka Y, Yoshida M. HFD-induced hepatic lipid accumulation and inflammation are decreased in Factor D deficient mouse. Sci. Rep. 2020;10:17593.

7. Horn P, Tacke F. Metabolic reprogramming in liver fibrosis. Cell Metab. 2024;36:1439–1455.

8. Fujiwara S, Izawa T, Mori M, Atarashi M, Yamate J, Kuwamura M. Dietary iron overload enhances Western diet induced hepatic inflammation and alters lipid metabolism in rats sharing similarity with human DIOS. Sci. Rep. 2022;12:21414.

9. Wang X, Zhang L, Dong B. Molecular mechanisms in MASLD/MASH-related HCC. Hepatology. 2024;

10. Loomba R, Friedman SL, Shulman GI. Mechanisms and disease consequences of nonalcoholic fatty liver disease. Cell. 2021;184:2537–2564.

11. Kew MC. Hepatic iron overload and hepatocellular carcinoma. Liver Cancer. 2014;3:31–40.

12. Tang D, Chen X, Kang R, Kroemer G. Ferroptosis: molecular mechanisms and health implications. Cell Res. 2021;31:107–125.

13. Bu L, Zhang Z, Chen J, Fan Y, Guo J, Su Y, et al. High-fat diet promotes liver tumorigenesis via palmitoylation and activation of AKT. Gut. 2024;73:1156–1168.

14. Furutani T, Hino K, Okuda M, Gondo T, Nishina S, Kitase A, et al. Hepatic Iron Overload Induces Hepatocellular Carcinoma in Transgenic Mice Expressing the Hepatitis C Virus Polyprotein. Gastroenterology. 2006;130:2087–2098.

15. Zhou Z-Q, Manguino D, Kewitt K, Intano GW, McMahan CA, Herbert DC, et al. Spontaneous hepatocellular carcinoma is reduced in transgenic mice overexpressing human O - methylguanine-DNA methyltransferase. Proceedings of the National Academy of Sciences. 2001;98:12566–12571.

16. Zavadil JA, Herzig MCS, Hildreth K, Foroushani A, Boswell W, Walter R, et al. C3HeB/FeJ Mice mimic many aspects of gene expression and pathobiological features of human hepatocellular carcinoma. Mol. Carcinog. 2019;58:309–320.

17. Men H, Young JL, Zhou W, Zhang H, Wang X, Xu J, et al. Early-Life Exposure to Low-Dose Cadmium Accelerates Diethylnitrosamine and Diet-Induced Liver Cancer. Oxid. Med. Cell. Longev. 2021;2021.

18. Zheng G, Bouamar H, Cserhati M, Zeballos CR, Mehta I, Zare H, et al. Integrin alpha 6 is upregulated and drives hepatocellular carcinoma progression through integrin α6β4 complex. Int. J. Cancer. 2022;151:930–943.

19. Sun X, Seidman JS, Zhao P, Troutman TD, Spann NJ, Que X, et al. Neutralization of Oxidized Phospholipids Ameliorates Non-alcoholic Steatohepatitis. Cell Metab. 2020;31:189–206.e8.

20. Kim D, Pertea G, Trapnell C, Pimentel H, Kelley R, Salzberg SL. TopHat2: accurate alignment of transcriptomes in the presence of insertions, deletions and gene fusions. Genome Biol. 2013;14:R36.

21. Li B, Dewey CN. RSEM: accurate transcript quantification from RNA-Seq data with or without a reference genome. BMC Bioinformatics. 2011;12:323.

22. Love MI, Huber W, Anders S. Moderated estimation of fold change and dispersion for RNA-seq data with DESeq2. Genome Biol. 2014;15:550.

23. Sherman BT, Hao M, Qiu J, Jiao X, Baseler MW, Lane HC, et al. DAVID: a web server for functional enrichment analysis and functional annotation of gene lists (2021 update). Nucleic Acids Res. 2022;50:W216–W221.

24. Yu G, Wang L-G, Han Y, He Q-Y. clusterProfiler: an R Package for Comparing Biological Themes Among Gene Clusters. OMICS. 2012;16:284–287.

25. Liberzon A, Birger C, Thorvaldsdóttir H, Ghandi M, Mesirov JP, Tamayo P. The Molecular Signatures Database Hallmark Gene Set Collection. Cell Syst. 2015;1:417–425.

26. Kolmykov S, Yevshin I, Kulyashov M, Sharipov R, Kondrakhin Y, Makeev VJ, et al. GTRD: an integrated view of transcription regulation. Nucleic Acids Res. 2021;49:D104–D111.

27. Barker M, Rayens W. Partial least squares for discrimination. J. Chemom. 2003;17:166–173.

28. Thévenot EA, Roux A, Xu Y, Ezan E, Junot C. Analysis of the Human Adult Urinary Metabolome Variations with Age, Body Mass Index, and Gender by Implementing a Comprehensive Workflow for Univariate and OPLS Statistical Analyses. J. Proteome Res. 2015;14:3322–3335.

29. Cui Q, Jiang T, Xie X, Wang H, Qian L, Cheng Y, et al. S-nitrosylation attenuates pregnane X receptor hyperactivity and acetaminophen-induced liver injury. JCI Insight. 2024;9.

30. Kabil O, Vitvitsky V, Banerjee R. Sulfur as a signaling nutrient through hydrogen sulfide. Annu. Rev. Nutr. 2014;34:171–205.

31. Visekruna A, Volkov A, Steinhoff U. A key role for NF-κB transcription factor c-Rel in T-lymphocyte-differentiation and effector functions. Clin. Dev. Immunol. 2012;2012:239368.

32. Pahl HL. Activators and target genes of Rel/NF-κB transcription factors. Oncogene. 1999;18:6853–6866.

33. Das D, Wang X, Chiu Y-C, Bouamar H, Sharkey FE, Lopera JE, et al. Integrative multi-omics characterization of hepatocellular carcinoma in Hispanic patients. JNCI: Journal of the National Cancer Institute. 2024;116:1961–1978.

34. Gardenghi S, Ramos P, Marongiu MF, Melchiori L, Breda L, Guy E, et al. Hepcidin as a therapeutic tool to limit iron overload and improve anemia in β-thalassemic mice. J. Clin. Invest. 2010;120:4466–77.

35. Yilmaz D, Tharehalli U, Paganoni R, Knoop P, Gruber A, Chen Y, et al. Iron metabolism in a mouse model of hepatocellular carcinoma. Sci. Rep. 2025;15:2180.

36. Zhou Q, Meng Y, Li D, Yao L, Le J, Liu Y, et al. Ferroptosis in cancer: From molecular mechanisms to therapeutic strategies. Signal Transduct. Target. Ther. 2024;9:55.

37. Luo Y, Niu G, Yi H, Li Q, Wu Z, Wang J, et al. Nanomedicine promotes ferroptosis to inhibit tumour proliferation in vivo. Redox Biol. 2021;42:101908.

38. Liu GY, Sabatini DM. mTOR at the nexus of nutrition, growth, ageing and disease. Nat. Rev. Mol. Cell Biol. 2020;21:183–203.

39. Mohamed NM, Mohamed RH, Kennedy JF, Elhefnawi MM, Hamdy NM. A comprehensive review and in silico analysis of the role of survivin (BIRC5) in hepatocellular carcinoma hallmarks: A step toward precision. Int. J. Biol. Macromol. 2025;311:143616.

40. Broadfield LA, Pane AA, Talebi A, Swinnen J V., Fendt S-M. Lipid metabolism in cancer: New perspectives and emerging mechanisms. Dev. Cell. 2021;56:1363–1393.

41. Que X, Hung M-Y, Yeang C, Gonen A, Prohaska TA, Sun X, et al. Oxidized phospholipids are proinflammatory and proatherogenic in hypercholesterolaemic mice. Nature. 2018;558:301–306.

42. Sun X, Seidman JS, Zhao P, Troutman TD, Spann NJ, Que X, et al. Neutralization of Oxidized Phospholipids Ameliorates Non-alcoholic Steatohepatitis. Cell Metab. 2020;31:189–206.e8.

43. Cho YK, Yoon YC, Im H, Son Y, Kim M, Saha A, et al. Adipocyte lysoplasmalogenase TMEM86A regulates plasmalogen homeostasis and protein kinase A-dependent energy metabolism. Nat. Commun. 2022;13:4084.

44. van Wouw SAE, van den Berg M, El Ouraoui M, Meurs A, Kingma J, Ottenhoff R, et al. Sterol-regulated transmembrane protein TMEM86a couples LXR signaling to regulation of lysoplasmalogens in macrophages. J. Lipid Res. 2023;64:100325.

45. Chen Z, Lu T, Huang L, Wang Z, Yan Z, Guan Y, et al. Circular RNA cia-MAF drives self-renewal and metastasis of liver tumor-initiating cells via transcription factor MAFF. Journal of Clinical Investigation. 2021;131.

46. Yu H, Cui Y, Guo F, Zhu Y, Zhang X, Shang D, et al. Vanin1 (VNN1) in chronic diseases: Future directions for targeted therapy. Eur. J. Pharmacol. 2024;962:176220.

47. Kang G, Han H-S, Koo S-H. NFIL3 is a negative regulator of hepatic gluconeogenesis. Metabolism. 2017;77:13–22.

48. Patsis C, Glyka V, Yiotakis I, Fragoulis EG, Scorilas A. l-DOPA Decarboxylase (DDC) Expression Status as a Novel Molecular Tumor Marker for Diagnostic and Prognostic Purposes in Laryngeal Cancer. Transl. Oncol. 2012;5:288–296.

49. He W, Wang X, Chen M, Li C, Chen W, Pan L, et al. Metformin reduces hepatocarcinogenesis by inducing downregulation of Cyp26a1 and CD8+ T cells. Clin. Transl. Med. 2023;13:e1465.

50. Kliewer SA, Goodwin B, Willson TM. The Nuclear Pregnane X Receptor: A Key Regulator of Xenobiotic Metabolism. Endocr. Rev. 2002;23:687–702.

51. Fujii M, Shibazaki Y, Wakamatsu K, Honda Y, Kawauchi Y, Suzuki K, et al. A murine model for non-alcoholic steatohepatitis showing evidence of association between diabetes and hepatocellular carcinoma. Med. Mol. Morphol. 2013;46:141–152.

52. Machado MV, Diehl AM. Pathogenesis of Nonalcoholic Steatohepatitis. Gastroenterology. 2016;150:1769–1777.

53. Gao D, Zhou Z, Chen L, Zheng J, Yang J. CGREF1 facilitates the cell proliferation, migration and invasion of hepatocellular carcinoma cells via regulation of EIF3H/ Wnt/β-Catenin signaling axis. BMC Cancer. 2025;25:435.

54. Wu S, Guo X, Zhou J, Zhu X, Chen H, Zhang K, et al. High expression of UNC5B enhances tumor proliferation, increases metastasis, and worsens prognosis in breast cancer. Aging. 2020;12:17079–17098.

55. Zhu R-X, Cheng ASL, Chan HLY, Yang D-Y, Seto W-K. Growth arrest-specific gene 2 suppresses hepatocarcinogenesis by intervention of cell cycle and p53-dependent apoptosis. World J. Gastroenterol. 2019;25:4715–4726.

56. Wang J, Wang T, Zhang Y, Liu J, Song J, Han Y, et al. CPEB1 enhances erastin-induced ferroptosis in gastric cancer cells by suppressing twist1 expression. IUBMB Life. 2021;73:1180–1190.

57. Encarnacion J, Smith DM, Choi J, Scafidi J, Wolfgang MJ. Activating transcription factor 3 regulates hepatic apolipoprotein A4 upon metabolic stress. J. Biol. Chem. 2025;301:108468.

58. Upadhyay G. Emerging Role of Lymphocyte Antigen-6 Family of Genes in Cancer and Immune Cells. Front. Immunol. 2019;10:819.

59. Fujimoto A, Furuta M, Shiraishi Y, Gotoh K, Kawakami Y, Arihiro K, et al. Whole-genome mutational landscape of liver cancers displaying biliary phenotype reveals hepatitis impact and molecular diversity. Nat. Commun. 2015;6:6120.

60. Liu Q, Zhang X, Qi J, Tian X, Dovjak E, Zhang J, et al. Comprehensive profiling of lipid metabolic reprogramming expands precision medicine for HCC. Hepatology. 2025;81:1164–1180.

61. Gao Q, Zhu H, Dong L, Shi W, Chen R, Song Z, et al. Integrated Proteogenomic Characterization of HBV-Related Hepatocellular Carcinoma. Cell. 2019;179:561–577.e22.

62. Pan Y, Li Y, Fan H, Cui H, Chen Z, Wang Y, et al. Roles of the peroxisome proliferator-activated receptors (PPARs) in the pathogenesis of hepatocellular carcinoma (HCC). Biomedicine & Pharmacotherapy. 2024;177:117089.

63. Wang M, Han J, Xing H, Zhang H, Li Z, Liang L, et al. Dysregulated fatty acid metabolism in hepatocellular carcinoma. Hepat. Oncol. 2016;3:241–251.

64. Liu Y, Zhu J, Jin Y, Sun Z, Wu X, Zhou H, et al. Disrupting bile acid metabolism by suppressing Fxr causes hepatocellular carcinoma induced by YAP activation. Nat. Commun. 2025;16:3583.

65. Xue C, Gu X, Zhao Y, Jia J, Zheng Q, Su Y, et al. Prediction of hepatocellular carcinoma prognosis and immunotherapeutic effects based on tryptophan metabolism-related genes. Cancer Cell Int. 2022;22:308.

66. Liu Y, Li G, Lu F, Guo Z, Cai S, Huo T. Excess iron intake induced liver injury: The role of gut-liver axis and therapeutic potential. Biomedicine & Pharmacotherapy. 2023;168:115728.

67. Raza S, Tewari A, Rajak S, Gupta P, Sinha RA. Extracellular RNA mediates iron-induced toxicity and inflammatory signalling in hepatic cells. Toxicol. Rep. 2025;14:102002.

68. Hilton C, Sabaratnam R, Drakesmith H, Karpe F. Iron, glucose and fat metabolism and obesity: an intertwined relationship. Int. J. Obes. 2023;47:554–563.

69. Todisco S, Convertini P, Iacobazzi V, Infantino V. TCA Cycle Rewiring as Emerging Metabolic Signature of Hepatocellular Carcinoma. Cancers (Basel). 2019;12.

70. Satapati S, Sunny NE, Kucejova B, Fu X, He TT, Méndez-Lucas A, et al. Elevated TCA cycle function in the pathology of diet-induced hepatic insulin resistance and fatty liver. J. Lipid Res. 2012;53:1080–1092.

71. Brahma MK, Gilglioni EH, Zhou L, Trépo E, Chen P, Gurzov EN. Oxidative stress in obesity-associated hepatocellular carcinoma: sources, signaling and therapeutic challenges. Oncogene. 2021;40:5155–5167.

72. McIntyre CL, Temesgen A, Lynch L. Diet, nutrient supply, and tumor immune responses. Trends Cancer. 2023;9:752–763.

