## Supplementary Figures for "Distinct and shared synergistic effects of high-fat and high-iron diets on liver tumorigenesis and transcriptomic remodeling in mice"

### Figure S1

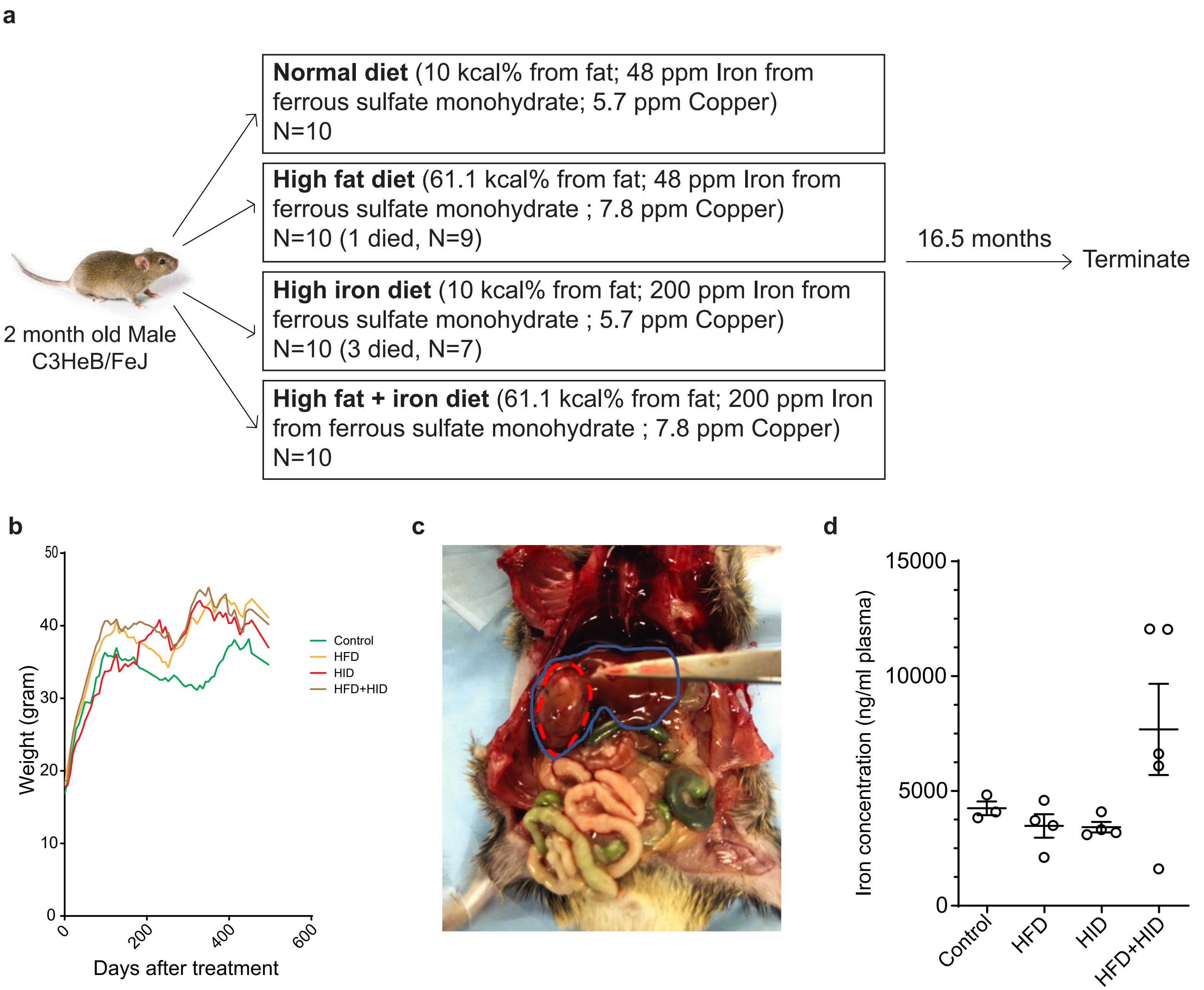


**Figure S1: Experimental design and pathophysiological impact of dietary interventions on mice.**
(a) Overview of the study design for dietary treatments in C3HeB/FeJ mice. Two-month-old male mice were assigned to one of four dietary groups: normal or control diet (Control, n=10), high-fat diet (HFD, n=10), high-iron diet (HID, n=10), and combined high-fat and high-iron diet (HFD+HID, n=10). Mice surviving until the end of the 16.5-month treatment period were euthanized, and their liver samples were collected for further analysis. (b) Body weight changes of mice across all dietary interventions during the treatment period.
(c) Representative image of a liver tumor in a C3HeB/FeJ mouse as indicated with the red oval circle.
(d) Plasma iron concentrations (ng/ml) across dietary groups.

#### Figure S2

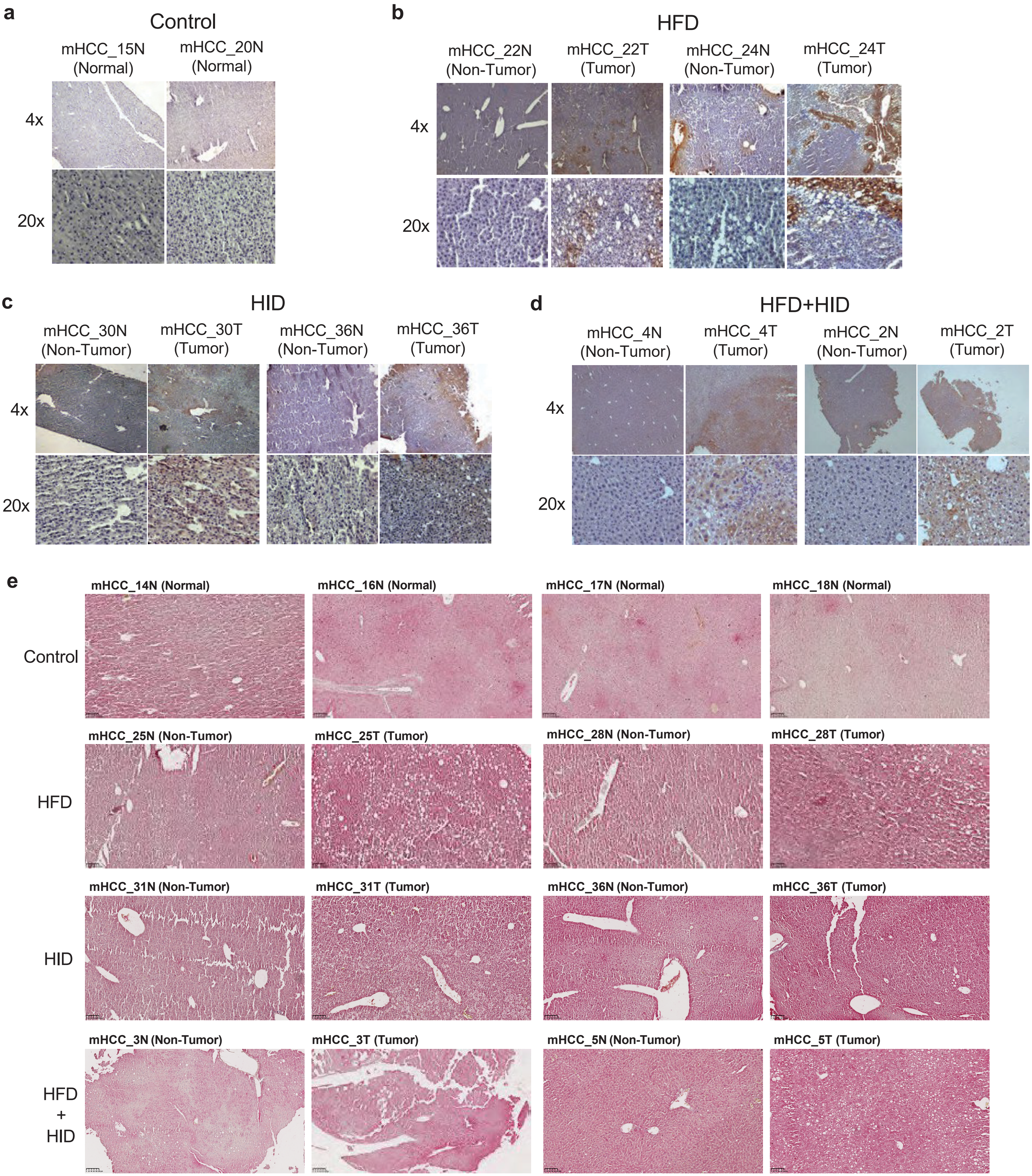


**Figure S2: Immunohistochemical analysis of phosphorylated S6-kinase (p-S6K) and OxPL levels in paraffin-embedded liver tissue sections of C3HeB/FeJ mice.** (a-d) Representative images of p-S6K immunostaining in liver tissues of mice treated with control diet (a), HFD (b), HID (c), and HFD+HID (d). The images were obtained with a 4X or 20X objective lens. (e) Representative images of liver sections (normal, non-tumor, and tumor tissues) stained with biotinylated IgM to assess oxidized phospholipid (OxPL) accumulation. Scale bar: 100 µm.

### Figure S3

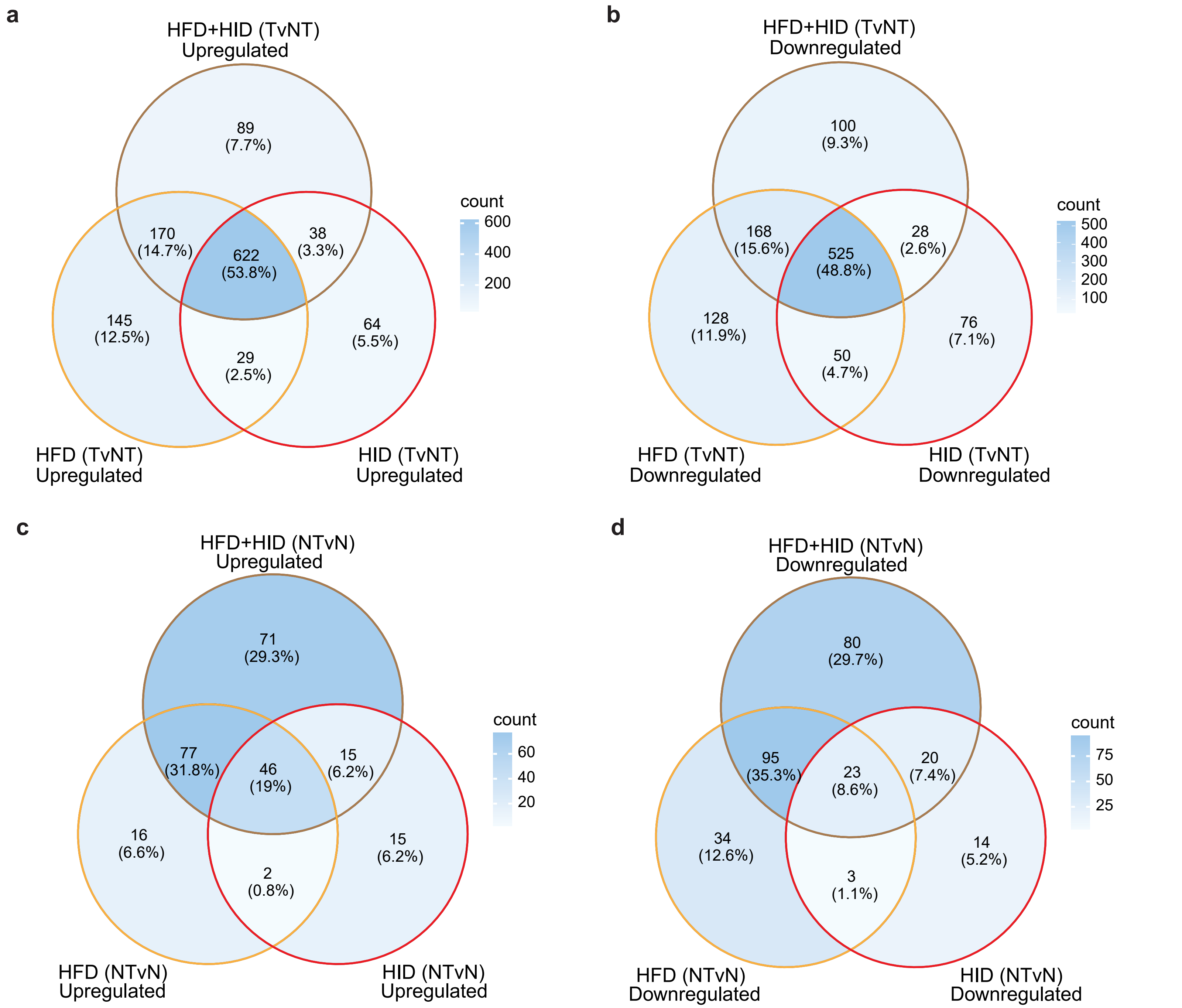


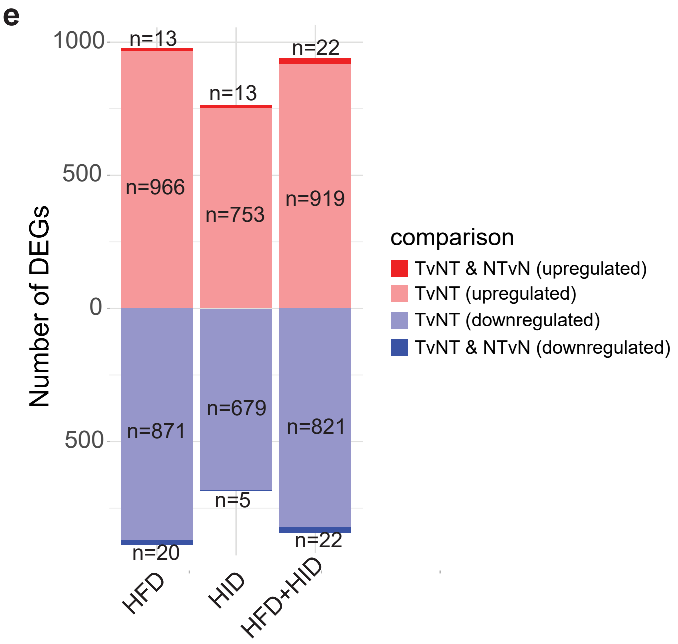


**Figure S3: Differential gene expression analysis across tumor, non-tumor, and normal liver tissues under different dietary conditions.** (a-b) Venn diagrams depicting the overlap of upregulated and downregulated differentially expressed genes (DEGs) (|Log2FC| > 1) in tumor versus paired non-tumor tissue comparisons (TvNT). (c-d) Venn diagrams illustrating the number of upregulated and downregulated DEGs in non-tumor liver tissues compared to normal liver tissues (NTvN) from control diet-treated mice across different dietary conditions. (e) Bar plot summarizing the number of DEGs unique to TvNT comparisons and those common between TvNT and NTvN comparisons.

#### Figure S4

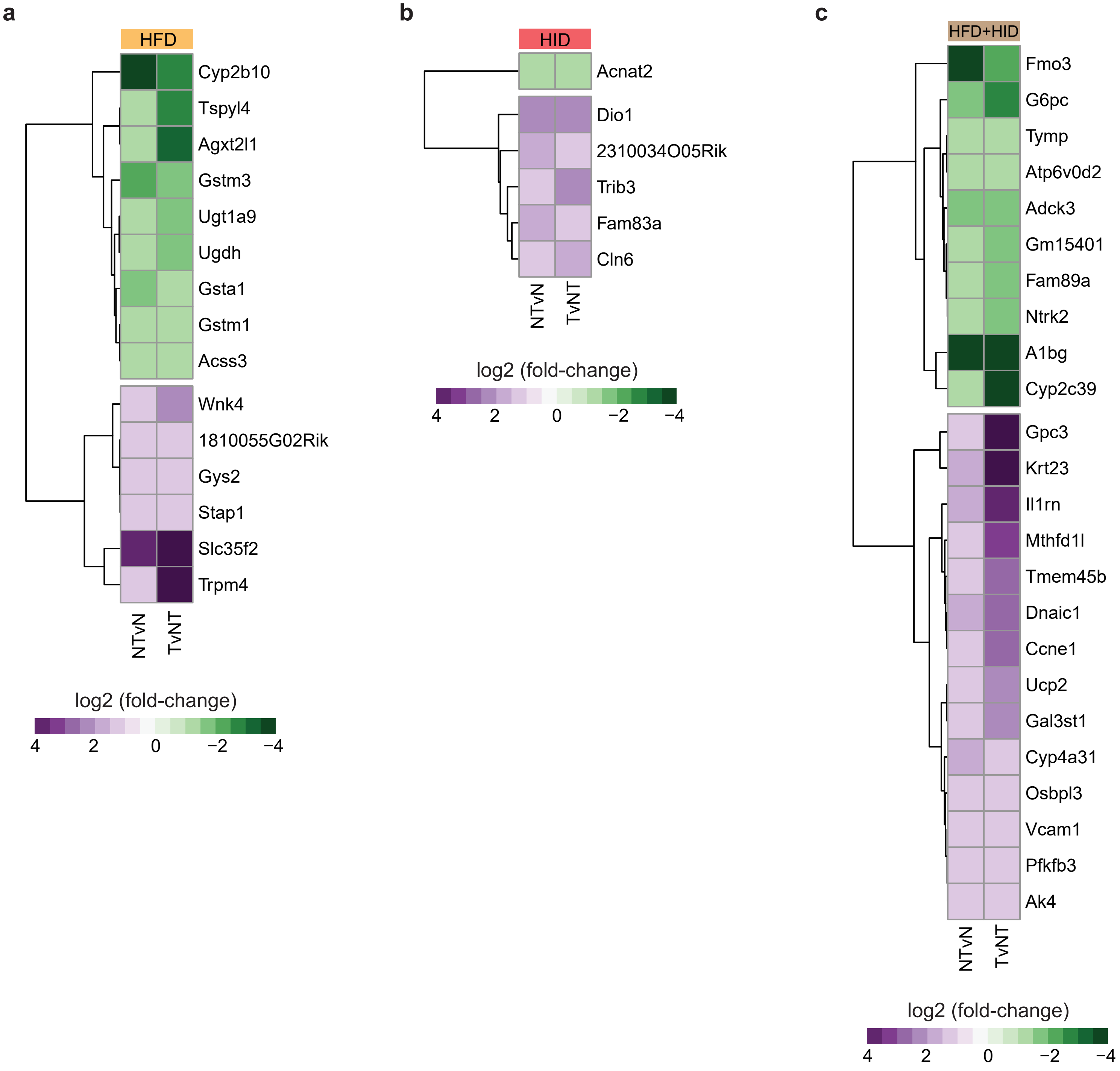


**Figure S4: Monotonic gene expression changes across liver tissue types in different dietary conditions.** List of DEGs showing monotonic upregulation or downregulation across normal liver, non-tumor, and tumor tissue samples, uniquely identified in mice treated with HFD (a), HID (b), and HFD+HID (c).

### Figure S5

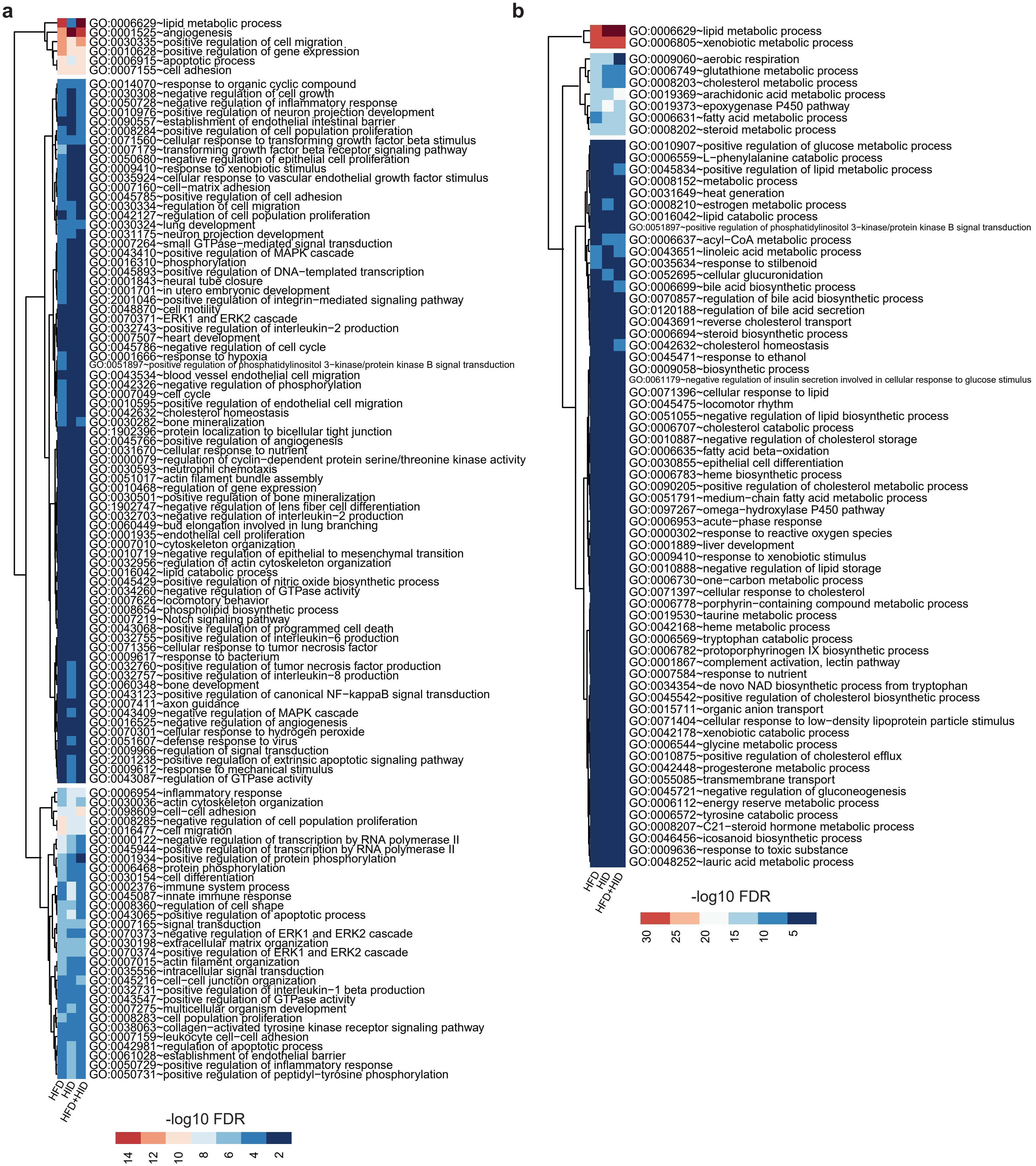


**Figure S5: Enrichment of gene ontology biological processes in tumor samples compared to paired non-tumor samples.** Gene Ontology Biological Process (GOBP) terms enriched in tumor samples relative to paired non-tumor samples based on differentially expressed genes (DEGs) identified as (a) upregulated and (b) downregulated regardless of dietary treatments. GOBP terms were clustered using Ward's method, based on the negative logarithm of false discovery rate (FDR) values obtained from enrichment analyses.

Figure S6

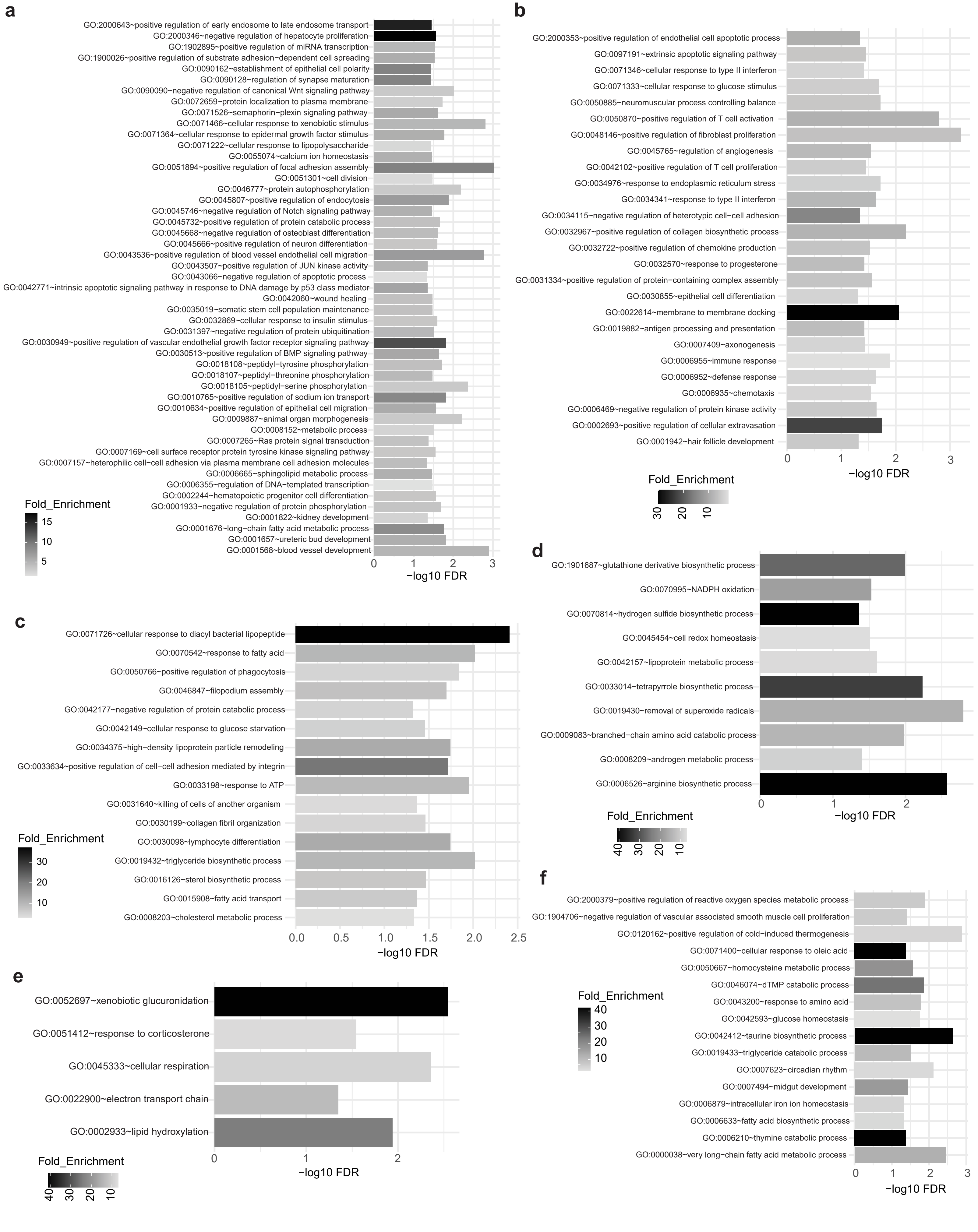

a
GO:2000643~positive regulation of early endosome to late endosome transport
GO:2000346~negative regulation of hepatocyte proliferation
GO:1902895~positive regulation of miRNA transcription
GO:1900026~positive regulation of substrate adhesion−dependent cell spreading
GO:0090162~establishment of epithelial cell polarity
GO:0090128~regulation of synapse maturation
GO:0090090~negative regulation of canonical Wnt signaling pathway
GO:0072659~protein localization to plasma membrane
GO:0071526~semaphorin−plexin signaling pathway
GO:0071466~cellular response to xenobiotic stimulus
GO:0071364~cellular response to epidermal growth factor stimulus
GO:0071222~cellular response to lipopolysaccharide
GO:0055074~calcium ion homeostasis
GO:0051894~positive regulation of focal adhesion assembly
GO:0051301~cell division
GO:0046777~protein autophosphorylation
GO:0045807~positive regulation of endocytosis
GO:0045746~negative regulation of Notch signaling pathway
GO:0045732~positive regulation of protein catabolic process
GO:0045668~negative regulation of osteoblast differentiation
GO:0045666~positive regulation of neuron differentiation
GO:0043536~positive regulation of blood vessel endothelial cell migration
GO:0043507~positive regulation of JUN kinase activity
GO:0043066~negative regulation of apoptotic process
GO:0042771~intrinsic apoptotic signaling pathway in response to DNA damage by p53 class mediator
GO:0042060~wound healing
GO:0035019~somatic stem cell population maintenance
GO:0032869~cellular response to insulin stimulus
GO:0031397~negative regulation of protein ubiquitination
GO:0030949~positive regulation of vascular endothelial growth factor receptor signaling pathway
GO:0030513~positive regulation of BMP signaling pathway
GO:0018108~peptidyl−tyrosine phosphorylation
GO:0018107~peptidyl−threonine phosphorylation
GO:0018105~peptidyl−serine phosphorylation
GO:0010765~positive regulation of sodium ion transport
GO:0010634~positive regulation of epithelial cell migration
GO:0009887~animal organ morphogenesis
GO:0008152~metabolic process
GO:0007265~Ras protein signal transduction
GO:0007169~cell surface receptor protein tyrosine kinase signaling pathway
GO:0007157~heterophilic cell−cell adhesion via plasma membrane cell adhesion molecules
GO:0006665~sphingolipid metabolic process
GO:0006355~regulation of DNA−templated transcription
GO:0002244~hematopoietic progenitor cell differentiation
GO:0001933~negative regulation of protein phosphorylation
GO:0001822~kidney development
GO:0001676~long−chain fatty acid metabolic process
GO:0001657~ureteric bud development
GO:0001568~blood vessel development
0
1
2
3
−log10 FDR
Fold_Enrichment
15
10
5
b
GO:2000353~positive regulation of endothelial cell apoptotic process
GO:0097191~extrinsic apoptotic signaling pathway
GO:0071346~cellular response to type II interferon
GO:0071333~cellular response to glucose stimulus
GO:0050885~neuromuscular process controlling balance
GO:0050870~positive regulation of T cell activation
GO:0048146~positive regulation of fibroblast proliferation
GO:0045765~regulation of angiogenesis
GO:0042102~positive regulation of T cell proliferation
GO:0034976~response to endoplasmic reticulum stress
GO:0034341~response to type II interferon
GO:0034115~negative regulation of heterotypic cell−cell adhesion
GO:0032967~positive regulation of collagen biosynthetic process
GO:0032722~positive regulation of chemokine production
GO:0032570~response to progesterone
GO:0031334~positive regulation of protein−containing complex assembly
GO:0030855~epithelial cell differentiation
GO:0022614~membrane to membrane docking
GO:0019882~antigen processing and presentation
GO:0007409~axonogenesis
GO:0006955~immune response
GO:0006952~defense response
GO:0006935~chemotaxis
GO:0006469~negative regulation of protein kinase activity
GO:0002693~positive regulation of cellular extravasation
GO:0001942~hair follicle development
0
1
2
3
−log10 FDR
Fold_Enrichment
30
20
10
c
GO:0071726~cellular response to diacyl bacterial lipopeptide
GO:0070542~response to fatty acid
GO:0050766~positive regulation of phagocytosis
GO:0046847~filopodium assembly
GO:0042177~negative regulation of protein catabolic process
GO:0042149~cellular response to glucose starvation
GO:0034375~high−density lipoprotein particle remodeling
GO:0033634~positive regulation of cell−cell adhesion mediated by integrin
GO:0033198~response to ATP
GO:0031640~killing of cells of another organism
GO:0030199~collagen fibril organization
GO:0030098~lymphocyte differentiation
GO:0019432~triglyceride biosynthetic process
GO:0016126~sterol biosynthetic process
GO:0015908~fatty acid transport
GO:0008203~cholesterol metabolic process
0.0
0.5
1.0
1.5
2.0
2.5
−log10 FDR
Fold_Enrichment
30
20
10
d
GO:1901687~glutathione derivative biosynthetic process
GO:0070995~NADPH oxidation
GO:0070814~hydrogen sulfide biosynthetic process
GO:0045454~cell redox homeostasis
GO:0042157~lipoprotein metabolic process
GO:0033014~tetrapyrrole biosynthetic process
GO:0019430~removal of superoxide radicals
GO:0009083~branched−chain amino acid catabolic process
GO:0008209~androgen metabolic process
GO:0006526~arginine biosynthetic process
0
1
2
−log10 FDR
Fold_Enrichment
40
30
20
10
e
GO:0052697~xenobiotic glucuronidation
GO:0051412~response to corticosterone
GO:0045333~cellular respiration
GO:0022900~electron transport chain
GO:0002933~lipid hydroxylation
0
1
2
−log10 FDR
Fold_Enrichment
40
30
20
10
f
GO:2000379~positive regulation of reactive oxygen species metabolic process
GO:1904706~negative regulation of vascular associated smooth muscle cell proliferation
GO:0120162~positive regulation of cold−induced thermogenesis
GO:0071400~cellular response to oleic acid
GO:0050667~homocysteine metabolic process
GO:0046074~dTMP catabolic process
GO:0043200~response to amino acid
GO:0042593~glucose homeostasis
GO:0042412~taurine biosynthetic process
GO:0019433~triglyceride catabolic process
GO:0007623~circadian rhythm
GO:0007494~midgut development
GO:0006879~intracellular iron ion homeostasis
GO:0006633~fatty acid biosynthetic process
GO:0006210~thymine catabolic process
GO:0000038~very long−chain fatty acid metabolic process
0
1
2
3
−log10 FDR
Fold_Enrichment
40
30
20
10

**Figure S6: Impact of dietary interventions on gene ontology biological process enrichment in HCC development.** Gene Ontology Biological Process (GOBP) terms enriched due to dietary-specific differentially expressed genes (DEGs) in tumors compared to non-tumor samples. GOBP terms significantly enriched from upregulated DEGs are shown for (a) high-fat diet (HFD-only, n = 49), (b) high-iron diet (HID-only, n = 26), and (c) combined high-fat and high-iron diet (HFD+HID-only, n = 16). GOBP terms significantly enriched from downregulated DEGs are presented for (d) HFD-only (n = 10), (e) HID-only (n = 5), and (f) HFD+HID-only (n = 16).

### Figure S7

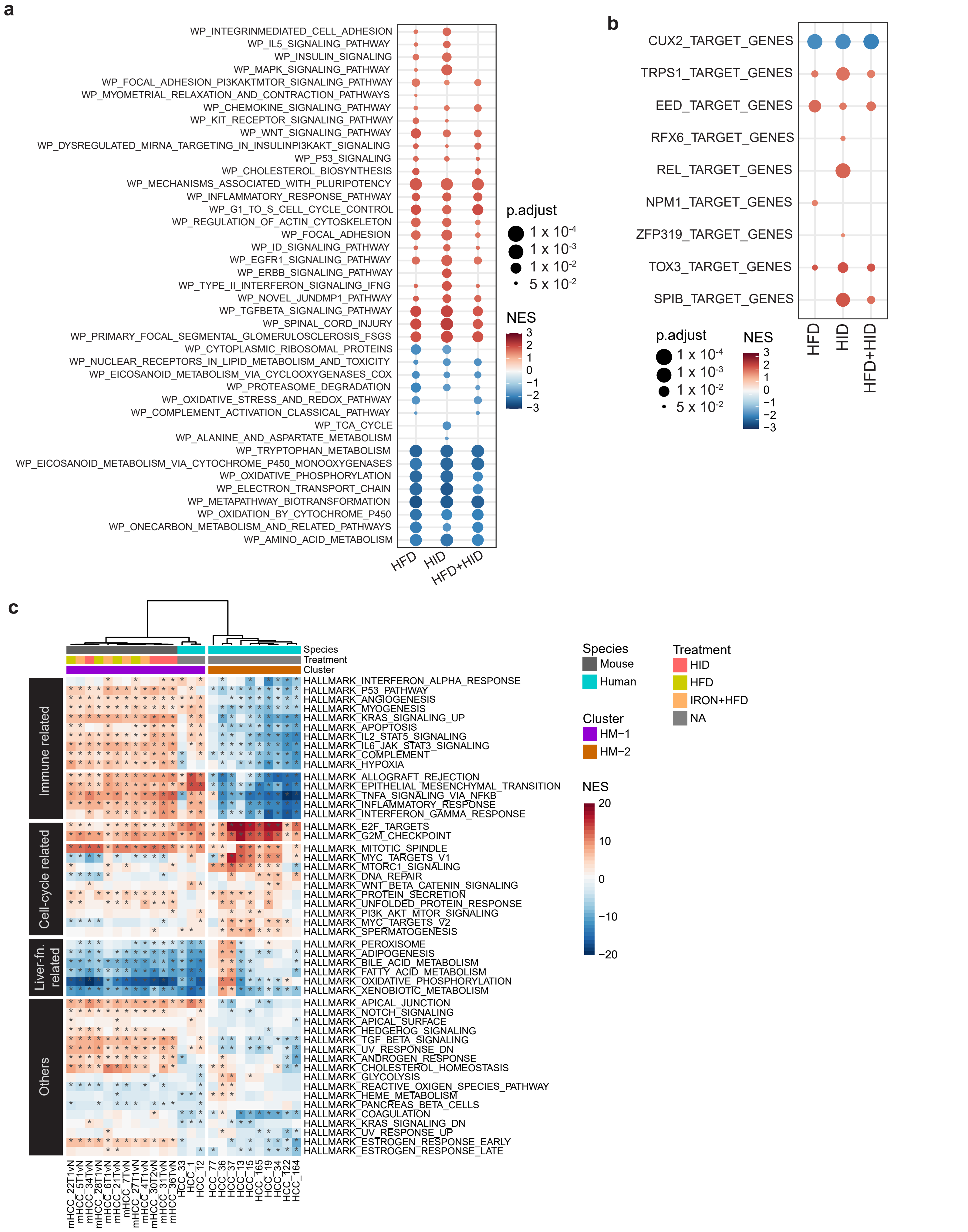
a
WP_INTEGRINMEDIATED_CELL_ADHESION
WP_IL5_SIGNALING_PATHWAY
WP_INSULIN_SIGNALING
WP_MAPK_SIGNALING_PATHWAY
WP_FOCAL_ADHESION_PI3KAKTMTOR_SIGNALING_PATHWAY
WP_MYOMETRIAL_RELAXATION_AND_CONTRACTION_PATHWAYS
WP_CHEMOKINE_SIGNALING_PATHWAY
WP_KIT_RECEPTOR_SIGNALING_PATHWAY
WP_WNT_SIGNALING_PATHWAY
WP_DYSREGULATED_MIRNA_TARGETING_IN_INSULINPI3KAKT_SIGNALING
WP_P53_SIGNALING
WP_CHOLESTEROL_BIOSYNTHESIS
WP_MECHANISMS_ASSOCIATED_WITH_PLURIPOTENCY
WP_INFLAMMATORY_RESPONSE_PATHWAY
WP_G1_TO_S_CELL_CYCLE_CONTROL
WP_REGULATION_OF_ACTIN_CYTOSKELETON
WP_FOCAL_ADHESION
WP_ID_SIGNALING_PATHWAY
WP_EGFR1_SIGNALING_PATHWAY
WP_ERBB_SIGNALING_PATHWAY
WP_TYPE_II_INTERFERON_SIGNALING_IFNG
WP_NOVEL_JUNDMP1_PATHWAY
WP_TGFBETA_SIGNALING_PATHWAY
WP_SPINAL_CORD_INJURY
WP_PRIMARY_FOCAL_SEGMENTAL_GLOMERULOSCLEROSIS_FSGS
WP_CYTOPLASMIC_RIBOSOMAL_PROTEINS
WP_NUCLEAR_RECEPTORS_IN_LIPID_METABOLISM_AND_TOXICITY
WP_EICOSANOID_METABOLISM_VIA_CYCLOOXYGENASES_COX
WP_PROTEASOME_DEGRADATION
WP_OXIDATIVE_STRESS_AND_REDOX_PATHWAY
WP_COMPLEMENT_ACTIVATION_CLASSICAL_PATHWAY
WP_TCA_CYCLE
WP_ALANINE_AND_ASPARTATE_METABOLISM
WP_TRYPTOPHAN_METABOLISM
WP_EICOSANOID_METABOLISM_VIA_CYTOCHROME_P450_MONOOXYGENASES
WP_OXIDATIVE_PHOSPHORYLATION
WP_ELECTRON_TRANSPORT_CHAIN
WP_METAPATHWAY_BIOTRANSFORMATION
WP_OXIDATION_BY_CYTOCHROME_P450
WP_ONECARBON_METABOLISM_AND_RELATED_PATHWAYS
WP_AMINO_ACID_METABOLISM
HFD
HID
HFD+HID
p.adjust
1 x 10-4
1 x 10-3
1 x 10-2
5 x 10-2
NES
3
2
1
0
−1
−2
−3
b
CUX2_TARGET_GENES
TRPS1_TARGET_GENES
EED_TARGET_GENES
RFX6_TARGET_GENES
REL_TARGET_GENES
NPM1_TARGET_GENES
ZFP319_TARGET_GENES
TOX3_TARGET_GENES
SPIB_TARGET_GENES
HFD
HID
HFD+HID
p.adjust
1 x 10-4
1 x 10-3
1 x 10-2
5 x 10-2
NES
3
2
1
0
−1
−2
−3
c
Species
Treatment
Cluster
Immune related
Cell-cycle related
Liver-fn. related
Others
HALLMARK_INTERFERON_ALPHA_RESPONSE
HALLMARK_P53_PATHWAY
HALLMARK_ANGIOGENESIS
HALLMARK_MYOGENESIS
HALLMARK_KRAS_SIGNALING_UP
HALLMARK_APOPTOSIS
HALLMARK_IL2_STAT5_SIGNALING
HALLMARK_IL6_JAK_STAT3_SIGNALING
HALLMARK_COMPLEMENT
HALLMARK_HYPOXIA
HALLMARK_ALLOGRAFT_REJECTION
HALLMARK_EPITHELIAL_MESENCHYMAL_TRANSITION
HALLMARK_TNFA_SIGNALING_VIA_NFKB
HALLMARK_INFLAMMATORY_RESPONSE
HALLMARK_INTERFERON_GAMMA_RESPONSE
HALLMARK_E2F_TARGETS
HALLMARK_G2M_CHECKPOINT
HALLMARK_MITOTIC_SPINDLE
HALLMARK_MYC_TARGETS_V1
HALLMARK_MTORC1_SIGNALING
HALLMARK_DNA_REPAIR
HALLMARK_WNT_BETA_CATENIN_SIGNALING
HALLMARK_PROTEIN_SECRETION
HALLMARK_UNFOLDED_PROTEIN_RESPONSE
HALLMARK_PI3K_AKT_MTOR_SIGNALING
HALLMARK_MYC_TARGETS_V2
HALLMARK_SPERMATOGENESIS
HALLMARK_PEROXISOME
HALLMARK_ADIPOGENESIS
HALLMARK_BILE_ACID_METABOLISM
HALLMARK_FATTY_ACID_METABOLISM
HALLMARK_OXIDATIVE_PHOSPHORYLATION
HALLMARK_XENOBIOTIC_METABOLISM
HALLMARK_APICAL_JUNCTION
HALLMARK_NOTCH_SIGNALING
HALLMARK_APICAL_SURFACE
HALLMARK_HEDGEHOG_SIGNALING
HALLMARK_TGF_BETA_SIGNALING
HALLMARK_UV_RESPONSE_DN
HALLMARK_ANDROGEN_RESPONSE
HALLMARK_CHOLESTEROL_HOMEOSTASIS
HALLMARK_GLYCOLYSIS
HALLMARK_REACTIVE_OXIGEN_SPECIES_PATHWAY
HALLMARK_HEME_METABOLISM
HALLMARK_PANCREAS_BETA_CELLS
HALLMARK_COAGULATION
HALLMARK_KRAS_SIGNALING_DN
HALLMARK_UV_RESPONSE_UP
HALLMARK_ESTROGEN_RESPONSE_EARLY
HALLMARK_ESTROGEN_RESPONSE_LATE
mHCC_22T1vN
mHCC_5T1vN
mHCC_34TvN
mHCC_28T1vN
mHCC_6T1vN
mHCC_21TvN
mHCC_7TvN
mHCC_27T1vN
mHCC_4T1vN
mHCC_30T2vN
mHCC_31TvN
mHCC_36TvN
HCC_33
HCC_1
HCC_12
HCC_77
HCC_36
HCC_37
HCC_13
HCC_15
HCC_165
HCC_19
HCC_34
HCC_122
HCC_164
Species
Mouse
Human
Cluster
HM−1
HM−2
NES
20
10
0
−10
−20
Treatment
HID
HFD
IRON+HFD
NA

**Figure S7: Gene set enrichment analysis across diet- and species-specific hepatocellular carcinoma.** (a) Gene set enrichment analysis (GSEA) highlights significantly enriched WikiPathways, with a focus on metabolic and signaling pathways under high-fat diet (HFD), high-iron diet (HID), and combined high-fat and high-iron diet (HFD+HID) conditions. (b) Enrichment analysis of transcription factor target gene sets reveals transcriptional regulators influenced by diet-induced changes across different dietary treatments. (c) Heatmap illustrating hallmark gene set enrichment in hepatocellular carcinoma (HCC) tumors compared to paired non-tumor samples across clusters and species (mouse and human). Hallmark gene sets are grouped into immune-related, cell-cycle-related, liver-function-related, and other categories. Clustering based on normalized enrichment scores (NES) shows that C3HeB/FeJ mice exhibit an immunologically "hot" (HM-1) tumor, with positive enrichment of most immune-related hallmark gene sets, irrespective of dietary conditions. Significant enrichment (FDR < 0.05) is indicated by asterisks.
